# Asthma-associated variants induce *IL33* differential expression through a novel regulatory region

**DOI:** 10.1101/2020.09.09.290098

**Authors:** Ivy Aneas, Donna C. Decker, Chanie L. Howard, Débora R. Sobreira, Noboru J. Sakabe, Kelly M. Blaine, Michelle M. Stein, Cara L. Hrusch, Lindsey E. Montefiori, Juan Tena, Kevin M. Magnaye, Selene M. Clay, James E. Gern, Daniel J. Jackson, Matthew C. Altman, Edward T. Naureckas, Douglas K. Hogarth, Steven R. White, Jose Luis Gomez-Skarmeta, Nathan Schoetler, Carole Ober, Anne I. Sperling, Marcelo A. Nobrega

**Affiliations:** Department of Human Genetics, University of Chicago, Chicago, IL 60637, USA; Department of Medicine, Section of Pulmonary and Critical Care Medicine, University of Chicago, Chicago, IL 60637, USA; Centro Andaluz de Biología del Desarrollo (CSIC/UPO/JA), Universidad Pablo de Olavide, Seville 41013, Spain; Department of Pediatrics, University of Wisconsin School of Medicine and Public Health, Madison, WI 53726, USA; Division of Allergy and Infectious Diseases, Department of Medicine, University of Washington, Seattle, WA 98195, USA

## Abstract

Genome-wide association studies (GWAS) have implicated the *IL33* locus in asthma, but the underlying mechanisms remain unclear. Here, we identify a 5 kb region within the GWAS-defined segment that acts as a strong regulatory element *in vivo* and *in vitro.* Chromatin conformation capture showed that this 5 kb region loops to the *IL33* promoter, potentially regulating its expression. Supporting this notion, we show that genotype at an asthma-associated SNP, rs1888909, located within the 5 kb region, is associated with *IL33* gene expression in human airway epithelial cells and IL-33 protein expression in human plasma, potentially through differential binding of OCT-1 (POU2F1) to the asthma-risk allele. Our data demonstrate that asthma-associated variants at the *IL33* locus mediate allele-specific regulatory activity and *IL33* expression, providing a novel mechanism through which a regulatory SNP contributes to genetic risk of asthma.

## Introduction

Asthma is a common and chronic inflammatory disease of the airways, with significant contributions from both genetic and environmental factors. Genetic factors account for more than half of the overall disease liability^1^ and GWAS have discovered >60 loci contributing to asthma disease risk^2^, with most of the associated variants located in noncoding regions. Linking these noncoding variants to genes and understanding the mechanisms through which they impart disease risk remains an outstanding task for nearly all asthma GWAS loci.

Among the most highly replicated asthma loci are variants near the genes encoding the cytokine IL-33 on chromosome 9p24.1 and its receptor, ST2 (encoded by *IL1RL1*), on chromosome 2q12.1, highlighting the potential importance of this pathway in the genetic etiology of asthma. A crucial role for IL-33 in allergic airway inflammation and bronchial airway hyperresponsiveness has been known since its discovery in 2005^3^. Studies in individuals with asthma and in murine asthma models have identified elevated levels of IL-33 protein in both sera and tissues^4, 5^. This cytokine is a potent inducer of type 2 immune responses through its receptor ST2, and has been broadly implicated in other allergic and inflammatory conditions, such as atopic dermatitis, allergic rhinitis, and eosinophilic esophagitis^6, 7, 8^.

The single nucleotide polymorphisms (SNPs) associated with increased asthma risk at the *IL33* GWAS locus reside within a linkage disequilibrium (LD) block in a noncoding genomic segment located 2.3 kb upstream of th *IL33* gene. We therefore posited that variants in this region impact on *IL33* expression by altering cis-regulatory element(s) that control quantitative, spatial and/or temporal-specific gene expression.

Previous studies of complex diseases have shown how regulatory variants in promoters and enhancer elements lead to an increased risk of disease through altering the expression of nearby genes^9, 10, 11, 12, 13^. In contrast, other types of cis-regulatory elements, including repressors and insulators (also known as enhancer blocking elements), are less understood and characterized than enhancers, but are also likely to be functionally modified by regulatory variants^14^.

Here, we combined genetic fine-mapping using GWAS data sets combined with functional annotations from relevant tissues to dissect the asthma-associated region upstream of the *IL33* gene. We identified a regulatory element harboring SNPs that control *IL33* expression. Genotypes at rs1888909, a SNP within this regulatory element, are associated with *IL33* expression in ethnically diverse populations, as well as IL-33 plasma protein levels. Our study provides functional insights into the role of common regulatory variants at the *IL33* locus and illustrate how a causal SNP can exert phenotypic effects by modulating the function of regulatory elements that do not fit into standard definitions of enhancers, insulators, or repressors.

## Results

### Defining the *IL33* locus asthma-associated critical region

Variants at the *IL33* locus have been robustly associated with asthma in GWAS of ethnically-diverse populations^2, 15, 16, 17, 18, 19, 20^. We first used LD between the most significantly associated SNP in each GWAS (referred to here on as the lead SNP) and other SNPs to define the region harboring potentially causal variants at this locus (Fig. 1a). Five lead SNPs were reported among seven large GWAS, defining an LD block spanning 41 kb in European ancestry individuals (chr9: 6,172,380-6,213,468, hg19; Supplementary Fig. 1a), which included 21 additional SNPs in LD (*r*^2^ ≥ 0.80) with at least one of the five lead SNPs (26 total SNPs). Because LD tracts are longer in European genomes compared to African ancestry genomes, we also sought results of GWAS in African Americans to potentially narrow this region. Three multi-ancestry GWAS ^16, 18, 19^ included African Americans, but only one^16^ provided GWAS results separately by ancestry. In that study, the lead GWAS SNP rs1888909 differed from the lead SNPs in the European ancestry (rs1342326^15^, rs928413^17^, rs7848215^20^ and rs992969^2^) or combined multi-ancestry (rs2381416^16^ and rs992969^18,19^) GWAS. Because there was so little LD in this region in African Americans, we used an *r*^2^ ≥ 0.40 to define LD. SNPs in LD with rs1888909 at *r*^2^ ≥ 0.4 in African Americans defined a region of 20 kb (chr9: 6,188,124-6,209,099, hg19; Supplementary Fig. 1b and Supplementary Table 1) that excluded two lead SNPs (rs928413^17^ and rs7848215^20^).

**Figure 1.**
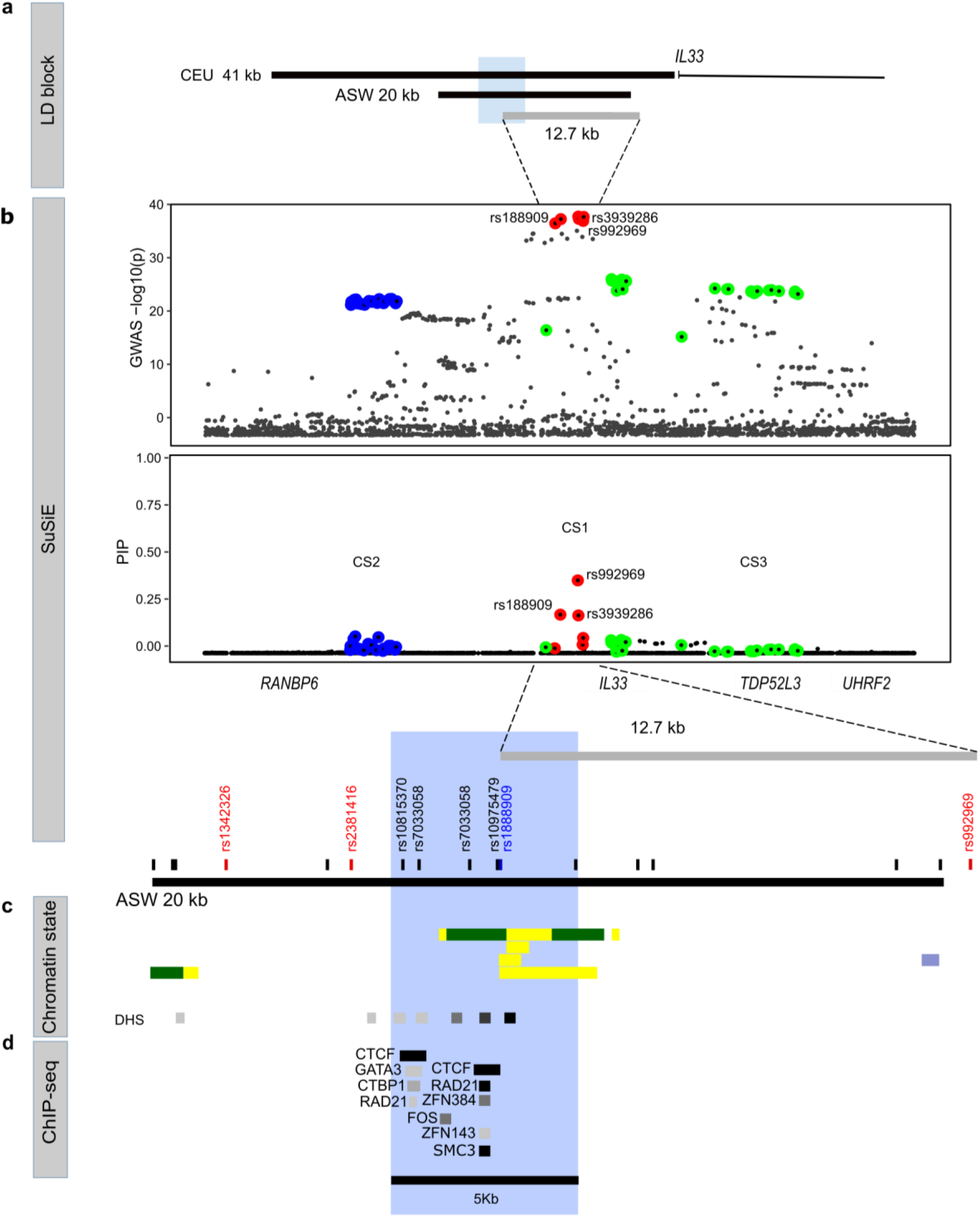
Epigenetic characterization of the asthma-associated critical region in the *IL33* locus. **a** Schematic organization of the *IL33* gene and the asthma-associated region (black bars) of European ancestry (CEU 41 kb, chr9: 6,172,380-6,213,468; hg19) and African ancestry (ASW 20 kb, chr9: 6,188,124-6,209,099; hg19) positioned upstream of exon 1. **b** Significance of SNP association in the GWAS2 (top) and fine mapping results (bottom) for each variant at the *IL33* locus (bottom). Colors indicate each credible set (CS) identified. CS1: red, CS2 blue; CS3 green. CS1 (rs992969, rs1888909, rs3939286) defines a region of 12.7 kb (chr9: 6,197,392 6,210,099; hg19). **c** Position of the lead GWAS SNPs (in red) and additional SNPs in high LD (r^2^≥0.8) with the lead SNPs (in black) within the ASW 20 kb LD region. The lead SNP rs1888909 in African ancestry is shown in blue. Chromatin states from Roadmap Epigenomics Project showing regions with potential regulatory activity. Yellow: active enhancer; green: transcribed sequence; blue: heterochromatin. DNase hypersensitive (DHS) sites indicating open chromatin regions are shown. Tissues (from the top): E096 Lung primary HMM; E095 Left ventricle primary HMM; E116 GM128781 Lymphoblastoid cell primary HMM, E122 HUVEC Umbilical Vein Endothelial Primary Cells Primary HMM **d** ChIP-seq data from ENCODE-3 cell lines (338 factors; 130 cell types) showing co-binding of CTCF, RAD2, ZFNs and SMC-3 at the 5 kb interval (blue shaded region; chr9: 6,194,500-6,199,500; hg19).

To gain more formal statistical support for this interval, we next used a Bayesian method, Sum of Single Effects (SuSiE)^21^, to fine-map the associated region and identify credible sets (CS) of SNPs at the *IL33* locus with high probabilities of being causal. Within each CS, variants are assigned posterior inclusion probabilities (PIPs), with higher PIPs reflecting higher probabilities of being causal and the sum of all PIPs within a CS always equaling 1. Using all SNPs at this locus from a GWAS of childhood onset asthma^2^ in British white subjects from the UK Biobank^22^, we identified three CSs (Fig.1b). CS1 contained variants with the highest PIPs and included six SNPs, all among those defined by LD with the lead GWAS SNPs, defining a 12.7 kb region approximately 22 kb upstream of the transcriptional start site of the *IL33* gene (chr9: 6,197,392-6,210,099; hg19) and overlapping with the regions defined by LD (Fig. 1a and Supplementary Table 2). The three CS1 SNPs with the highest PIPs included the lead SNPs in two multi-ancestry GWAS (rs992969; PIP 0.391), the lead SNP in the African American GWAS^16^ (rs1888909; PIP 0.209), and a SNP in LD with the lead SNPs (rs3939286; PIP 0.205). We note that two other CSs of 25 SNPs (CS2) and 28 SNPs (CS3) were identified by SuSiE, suggesting additional, independent regions potentially regulating the expression of *IL33* or other genes.

Next, we used Roadmap Epigenome^23^ and ENCODE^24^ data to annotate the regulatory landscape of the more inclusive 20 kb asthma-associated interval defined by LD. This segment was enriched with chromatin marks and DNase hypersensitive sites (Fig. 1c), suggestive of regulatory potential in multiple cell types. We also identified two CTCF sites within 2 kb of each other with evidence of CTCF binding in multiple cell lines (84 cell lines have CTCF binding to site 1 and 142 have CTCF binding to site 2 out 194 ENCODE-3 lines). In addition, the cohesin complex RAD21-SMC-3 subunits and zinc-finger proteins such as ZNF384 and ZNF143 also bind to this region (Fig. 1d). Binding of this multi-subunit complex along with CTCF can provide sequence specificity for chromatin looping to promoters or have insulator functions^25, 26^.

Interestingly, the three lead GWAS SNPs (rs1342326^15^, rs2381416^16^ and rs992969^2, 18, 19^) in the LD-defined 20 kb region did not overlap with regions of open chromatin or transcription factor binding; one, rs992969^2, 18, 19^, mapped within heterochromatin (thick blue horizontal bar) in virtually every ENCODE cell line (Supplementary Fig. 2). Heterochromatin is highly compacted region in the genome and not actively involved in gene regulation. For these reasons, it is not likely any of these three lead SNPs within the LD-defined region are causative of the asthma association. In contrast, five SNPs in high LD with the asthma-associated lead SNPs, including two from CS1 and including the lead African American GWAS SNP (rs1888909), overlapped with a region of open chromatin and CTCF, cohesin and ZNF binding, delineating a discrete 5 kb region (chr9: 6,194,500-6,199,500, hg19) (Fig. 1c-d). Collectively, these data reveal a 5 kb region that harbors both asthma-associated SNPs and marks of regulatory activity that could modulate *IL33* expression through long-range interactions.

### Regulatory properties of a 5 kb region upstream of *IL33*

Having identified a region of interest, we sought to determine its impact on *IL33* expression. Because *IL33* is expressed in multiple tissues, we used an *in vivo* model system to assay for the spatial regulatory properties of this 5 kb region. However, the lack of evolutionary conservation at the locus between human and mouse (Supplementary Fig. 3) required the creation of a “humanized” transgenic mouse model. For this, we used a human Bacterial Artificial Chromosome (BAC - clone RP11-725F15), approximately 166 kb long, spanning the coding region of the *IL33* gene and its upstream sequences, including the 20 kb asthma-associated region. We recombined a cassette containing a E2-Crimson reporter with a 3’ stop codon into exon 2, in frame with the *IL33* start codon (ATG). Any *IL33* regulatory regions within this BAC would drive E2-Crimson expression, mimicking *IL33* endogenous spatio-temporal expression patterns. Also, to directly test the regulatory impact of the 5 kb region, we selectively deleted this asthma-associated DNA segment from the full BAC and assessed the resultant *IL33* expression *in vivo*, in mice harboring the full length or the 5 kb deletion BACs (Fig. 2a).

**Figure 2.**
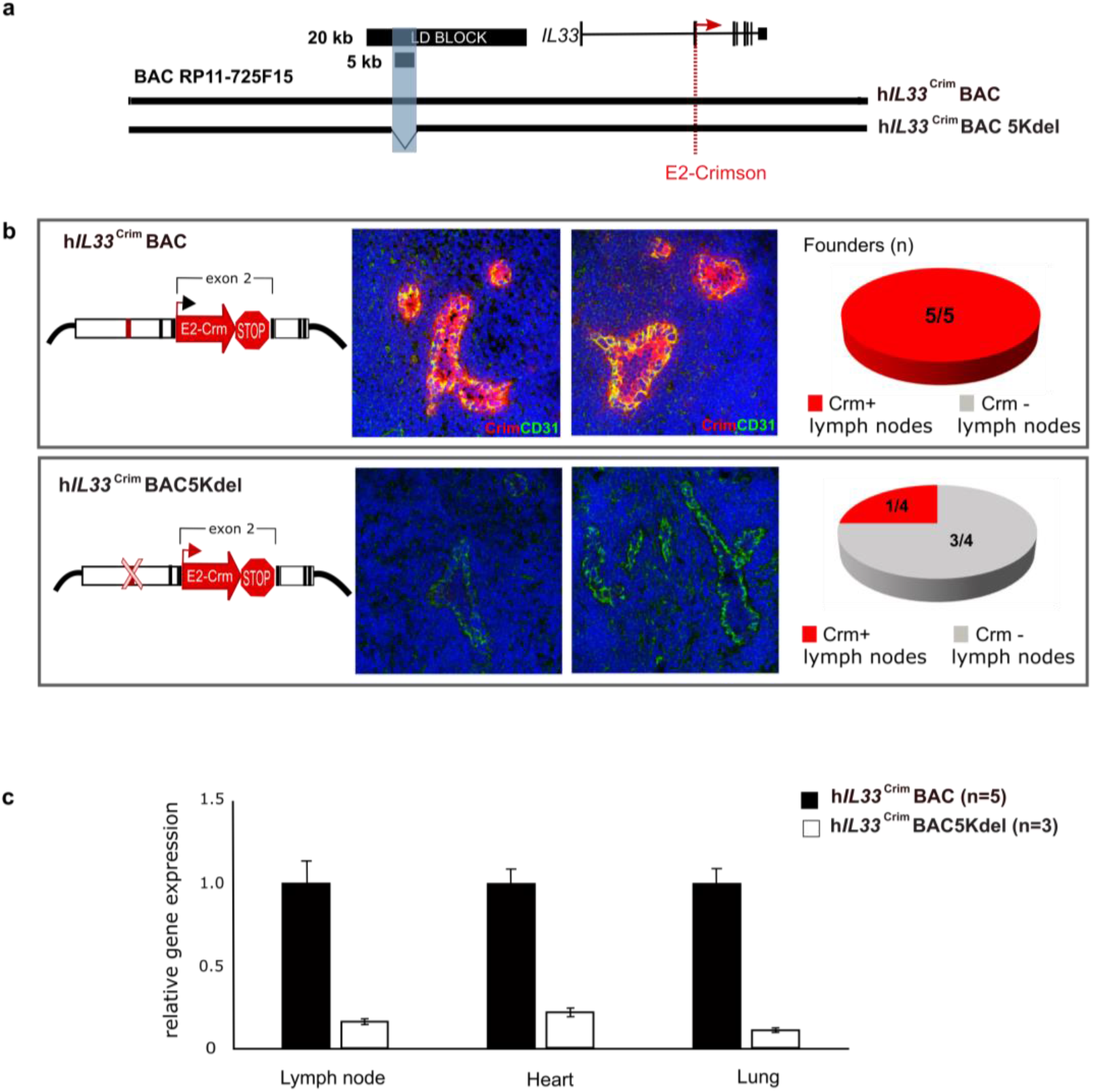
The *IL33-containing* BAC in transgenic mice encodes human-specific regulatory patterns and demonstrates the importance of the 5 kb noncoding segment for proper *IL33* expression. **a** Schematic of human BAC clone RP11-725F15 (166 kb) spanning the entire coding region of *IL33* and its upstream region including the 20 kb asthma-associated interval and the 5 kb region of interest shaded in blue (black bars). To produce a human *IL33* reporter strain, a cassette containing E2-Crimson with a stop sequence was inserted into exon 2, in frame with the *IL33* translational start site (red dotted line). Transgenic mice were generated with either the full BAC (hI*L33*^Crim^BAC) or a BAC containing a deletion of the 5 kb interval within the LD block (h*IL*33^Crim^ BAC5Kdel). **b** Immunofluorescence staining of mouse peripheral lymph node sections of E2-Crimson in h*IL33*^Crim^ BAC mice (upper panels) or h*IL33*^Crim^ BAC5Kdel (lower panels). Representative founder BAC transgenic mice are shown. Sections were stained with anti-E2-Crimson (red) and the mouse endothelial cell marker CD31 (green). Hoescht staining for nuclei is in blue. **c** qPCR analysis of E2-Crimson mRNA obtained from lymph node (LN), heart and lung in both BAC strains are shown

Immunofluorescence staining of peripheral lymph node in full BAC transgenic mice (h*IL33*^Crim^BAC) showed that E2-Crimson fluorescent protein is highly expressed in this tissue (Fig. 2b, upper panel and Supplementary Fig. 4a). Strikingly, constitutive expression of the E2-Crimson reporter was co-localized with the endothelial marker CD31 and observed in high endothelial venule (HEV) cells in mouse lymph nodes. This observation validates the species-specificity of the E2-Crimson expression, as previous studies report that while IL-33 is produced by HEV in humans, it is not found in murine HEV^**27**^. In contrast, deletion of the 5 kb region in the reporter BAC (h*IL33*^Crim^BAC5kdel) significantly depleted E2-Crimson immunostaining in lymph nodes compared to full BAC mice (Fig. 2b, lower panel and Supplementary Fig. 4b). The 5 kb deletion also significantly reduced E2-Crimson mRNA expression in heart and lung in 3 out of 4 independent lines (Fig. 2c). These results show that the BAC encodes human-specific regulatory patterns *in vivo* and demonstrate the importance of the 5 kb noncoding segment for proper *IL33* expression.

### Long-range chromatin interactions in the *IL33* locus

We next assessed the physical interactions between the asthma-associated region and the *IL33* promoters, by performing circular chromosome conformation capture followed by high-throughput sequencing (4C-seq) in cells obtained from h*IL33*^Crim^BAC and h*IL33*^Crim^BAC5kdel mice. Chromatin state annotations from Roadmap Epigenome suggest the presence of distal regulatory elements, such as enhancers (yellow bars) distributed across the human BAC region containing the *IL33* gene and its upstream non-coding DNA (Fig. 3a).

**Figure 3.**
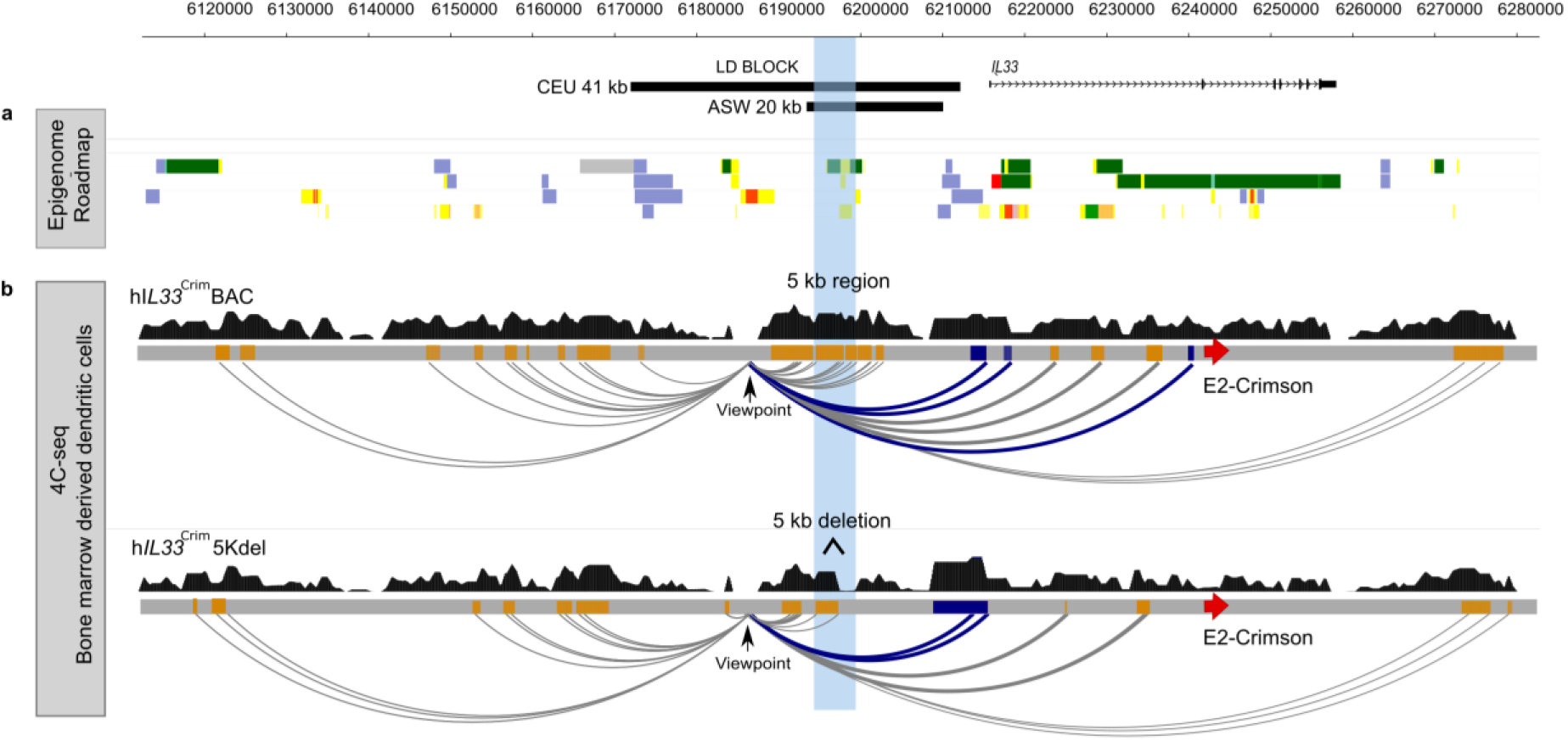
Interaction maps show looping of the asthma-associated region to the *IL33* promoter. **a** Roadmap Epigenome data from heart (E095 Left ventricle primary HMM), lung (E096 Lung primary HMM), LCL (E116 GM128781 Lymphoblastoid cell primary HMM) and endothelial cells (E122 HUVEC Umbilical Vein Endothelial Primary Cells Primary HM). Chromatin state assignments are indicated as: transcribed, green; active enhancer, yellow; active promoter, red; repressed, gray; heterochromatin, blue. **b** 4C-seq in bone marrow derived dendritic cells obtained from mice containing the BAC (h*IL33*^Crim^BAC, top) and a deletion of the 5 kb interval within the LD block (h*IL33*^Crim^ BAC5Kdel, bottom). Gray bars correspond to a schematic representation of the human BAC showing the E2-Crimson reporter insertion site on exon 2 of the *IL33* gene (red arrow) and the asthma associated 5 kb region (blue shaded box). In both experiments, reads were mapped to the coordinates corresponding to the human BAC (chr9: 6,112,733-6,279,294; hg19) and peaks shown in dark blue and orange represent the promoter and distal elements, respectively. Arcs depict interactions from the viewpoint located upstream the 5 kb region. Interactions between the LD region, near to the 5 kb region (viewpoint), and the *IL33* promoters are noted in blue

We positioned the viewpoint directly upstream of the 5 kb region in order to capture interactions between this genomic region and the entire adjacent *IL33* locus both in h*IL33*^Crim^BAC cells and h*IL33*^Crim^BAC5kdel cells (Fig. 3). In both cases, we observed interactions between the viewpoint and several potential regulatory regions located upstream and downstream (gray arcs), including the less frequently used promoter of the long transcript as well as the predominantly used promoter for the short transcript of *IL33* (blue arcs). Deletion of the 5 kb fragment reduced the interactions at the *IL33* locus compared to those observed using the full BAC, including the loss of the interaction with the short transcript promoter, confirming the necessity of this region for proper *IL33* regulation and explaining the loss of E2-Crimson expression in h*IL33*^Crim^BAC5kdel mice (Fig. 2b-c).

Because we are limited to measure interactions within the 166 kb of the human BAC in mouse cells, we cannot exclude the possibility that the 5 kb region also physically interacts with and regulates other genes in the topological associated domain (TAD) containing *IL33* in humans. To address this possibility, we carried out promoter capture Hi-C in human lymphoblastoid cell lines (LCLs). Our data show interactions between the asthma-associated LD block region and both *IL33* promoters but not to other genes (Supplementary Fig.5). These data further support the notion that the asthma-associated LD region harbors regulatory elements that physically interact with *IL33* to regulate its expression.

### Asthma-associated SNPs modify regulatory properties of the 5 kb region

To functionally characterize the regulatory impact of allelic variants of asthma-associated SNPs in the 5 kb region, we utilized a combination of *in vitro* and *in vivo* reporter assays. Because this region overlaps with chromatin states suggestive of enhancer function in some cell types, and because its deletion in the BAC resulted in a generalized loss of reporter expression *in vivo*, we hypothesized that the 5 kb region corresponds to an enhancer. To test the regulatory potential of this DNA segment *in vitro*, we cloned both the risk and non-risk haplotypes of the 5 kb fragment, as well as a shorter 1 kb fragment centered on the accessible chromatin segment from ENCODE (Fig. 1c) in a luciferase vector and transfected them in human cell lines. We used an immortalized human aortic endothelial cell line (TeloHAEC), as the ENCODE project chromatin data annotate this region as a putative enhancer in endothelial cells. We failed to detect enhancer activity of either haplotype in either the 5 kb or 1 kb fragments (Supplementary Fig. 6).

Interestingly, upon close inspection of ENCODE data we notice that this 5 kb region is not bound by transcription factors usually associated with enhancer activity, including transcriptional activators, repressors, or RNA PolI. Rather, it is bound by both CTCF and subunits of the cohesin complex across multiple cell lines (Fig. 1d). Both CTCF and cohesin are key determinants of chromatin loop formation and stabilization, including the positioning of regulatory elements close to the promoters of their target genes, or serving as insulators, with enhancer blocking properties.

To test the enhancer blocking properties of the 5 kb fragment, we first used an *in vitro* luciferase-based assay in which candidate sequences are cloned between a strong promoter (SV40) and a strong enhancer (HS2 element in human beta-globin LCR)^28^. Luciferase expression is driven by the enhancer and promoter elements, and a decrease in luciferase activity would be interpreted as enhancer blocking activity, with the enhancer not able to loop to the adjacent SV40 promoter. When the 5 kb region was cloned into this vector we observed prominent enhancer-blocking activity (Fig. 4a) and significant differences were also observed between fragments containing either the risk or non-risk alleles for the five SNPs, with the risk allele showing weakened enhancer blocking activity (Fig. 4b; p=0.001). As a control, we used a DNA sequence located 60 kb upstream of *IL33* and devoid of any epigenetic marks of active chromatin and with no evidence of CTCF or cohesin binding. This control sequence had no significant impact on the reporter gene expression (Fig. 4a).

**Figure 4.**
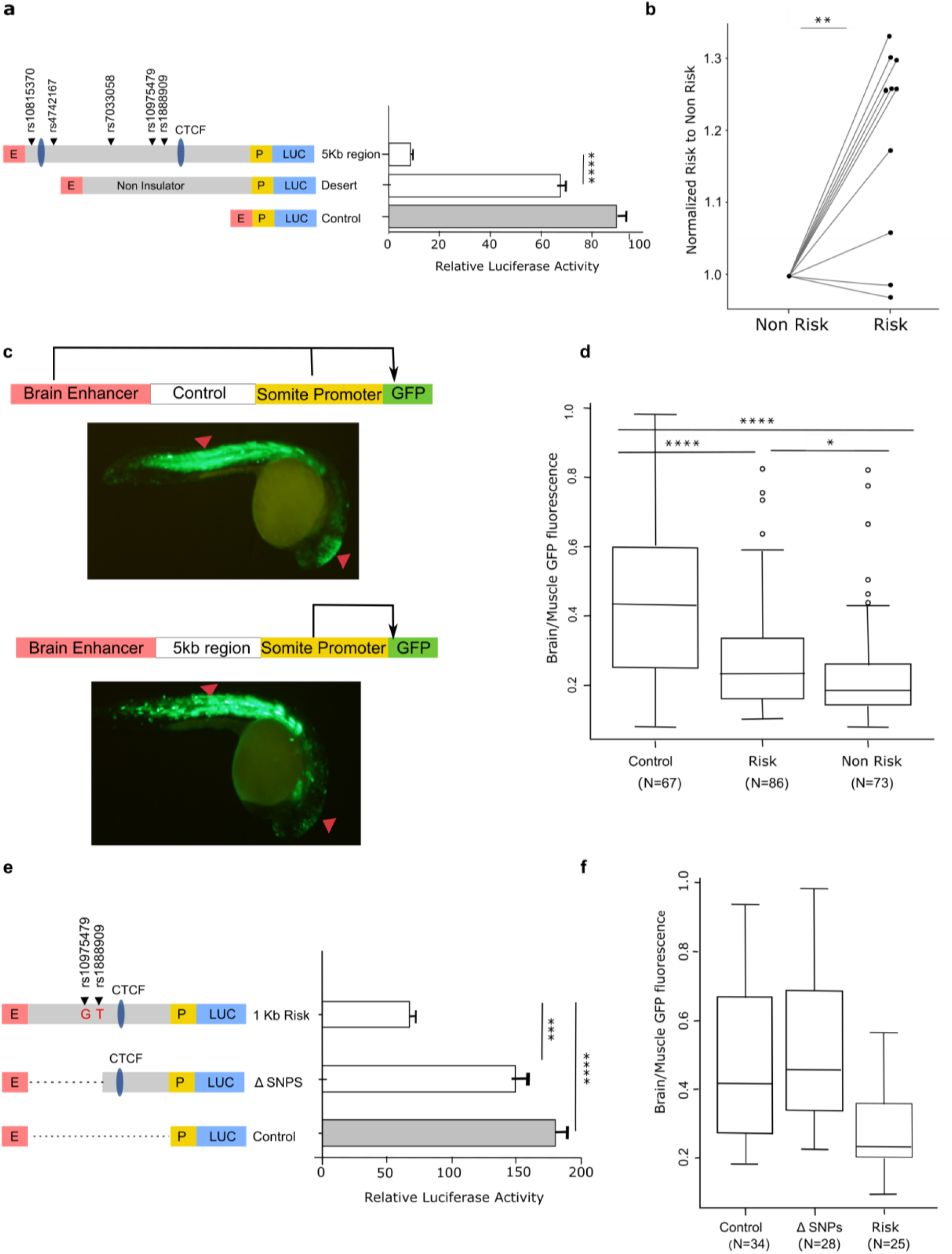
Impact of the asthma-associated variants in the regulatory property of the 5 kb region. **a** *In vitro* transgenic reporter assay. Luciferase based enhancer barrier assay using 5 kb constructs (chr9: 6,194,500-6,199,500; hg19) that were cloned between HS2 enhancer (E) and SV40 promoter (P) sequences. SNPs in the construct are noted (black arrowheads). Results are from 3 independent experiments. ****p<0.0001, unpaired t-test. **b** Luciferase activity values of the risk construct is shown as fold change over the activity obtained in the non-risk sequence. Results shown represent data from 10 independent experiments. **p=0.0014, two-way ANOVA. **c** *In vivo* zebrafish transgenic reporter assay. Green fluorescent protein (GFP) expression 24 hours post fertilization (hpf) in mosaic F0 embryos injected with vectors containing a control sequence (top panel) or 5 kb interval sequence (bottom panel). **d** Comparison between 5 kb constructs containing risk or non-risk alleles for enhancer blocking property. Data is presented as midbrain/somites EGFP intensity ratio compared with empty gateway vector which has no enhancer blocking activity. ****p=1.1e-5; * p=0.041, ANOVA pairwise T-test. **e** 1 kb DNA fragment (chr9: 6,197,000-6,197,917; hg19) containing the risk alleles for asthma variants rs10975479 and rs1888909 or a 400bp deletion in the region harboring those SNPs (chr9: 6,197,399-6,197,914; hg19) were assayed for luciferase assay in K562 cells. Results are representative of 3 independent experiments ***p=0.0005, unpaired t-test. **f** Zebrafish reporter assay comparing the 1 kb DNA fragment and fragment with the 400bp deletion (ΔSNP). Data are presented as in (d).

To test the this enhancer blocking property *in vivo* we used a zebrafish reporter assay^29^. The 5 kb region was cloned in a reporter cassette containing Green Fluorescent Protein (GFP) driven by a cardiac actin promoter and a midbrain enhancer. An enhancer blocking element cloned in this vector would restrict the access of the midbrain enhancer to the GFP reporter gene, while the ability of the actin promoter to activate GFP in skeletal muscle and somites would be maintained (Fig. 4c). In these experiments, the 5 kb region led to a decreased mid-brain-specific GFP signal when compared to the control sequence, which displayed no enhancer blocking activity (Fig. 4d; p<0.001). Similar to the data in the *in vitro* luciferase experiments, there was a significant difference in GFP activity between fragments containing the risk or non-risk alleles (p=0.041), with the risk alleles resulting in reduced enhancer blocking activity. Taken together, our data provide evidence that the 5 kb region has enhancer blocking properties *in vitro* and *in vivo*, and that alleles of asthma-associated SNPs within this region are able to modulate this property.

The 5 kb fragment contains two CTCF sites within 2 kb of each other and 5 asthma-associated SNPs. To determine the impact of the SNPs on its regulatory function, we cloned a smaller 1 kb fragment, including one CTCF site and the SNPs rs10975479 and rs1888909, located 15 bp apart from each other, and displaying the highest LD (r^2^) with the tag SNP rs992969 (0.56 and0.96, in CEU respectively). This segment of the 5 kb region overlaps with the minimal critical region predicted by fine-mapping by SuSie (Fig. 1a). This shorter fragment still showed prominent enhancer blocking activity *in vitro,* in luciferase assays (Fig. 4e). Furthermore, deletion of a 400 bp within the 1 kb element, harboring the SNPs rs10975479 and rs1888909, abrogated the enhancer blocking activity of this fragment. Importantly, this 400 bp deletion does not span the CTCF binding site within the 1 kb fragment. Testing these fragments in our *in vivo* zebrafish transgenic reporter assay confirmed the 1 kb fragment enhancer blocking activity and the functional importance of one or both SNPs within the 400 bp deleted region in regulating GFP expression (Fig. 4f). Together, these data characterize a regulatory region upstream of the *IL33* locus and implicate the asthma-associated SNPs rs10975479 and rs1888909 in regulating *IL33* expression.

### Differential binding of OCT-1 to allelic variants of rs1888909

Based on the regulatory properties of this region *in vitro* and *in vivo*, we hypothesized that the risk alleles of rs1888909 may alter transcription factor binding, resulting in altered regulatory activity and *IL33* expression. Although not predicted to be a causal SNP (Fig. 1b), we also studied rs10975479 (G/A) as it is only 15 bp away from rs1888909. We performed an electrophoretic mobility shift assay (EMSA) using small labeled DNA probes and unlabeled competitors spanning four different combinations of the risk or non-risk alleles for variants rs10975479 (G/A) and rs1888909 (T/C). Upon incubation with nuclear extract, we observed differences in binding patterns between the risk (G-T) and non-risk (A-C) probes, suggesting that different transcription factor binding complexes are formed in the presence of these two alleles (Fig. 5a). The major changes in binding are observed with both probes carrying the rs1888909 (T) allele, thereby indicating that the difference is driven by the risk allele T of this SNP. These data demonstrate that risk allele (T) of the asthma-associated SNP rs1888909 alters protein binding properties, possibly influencing *IL33* expression.

**Figure 5.**
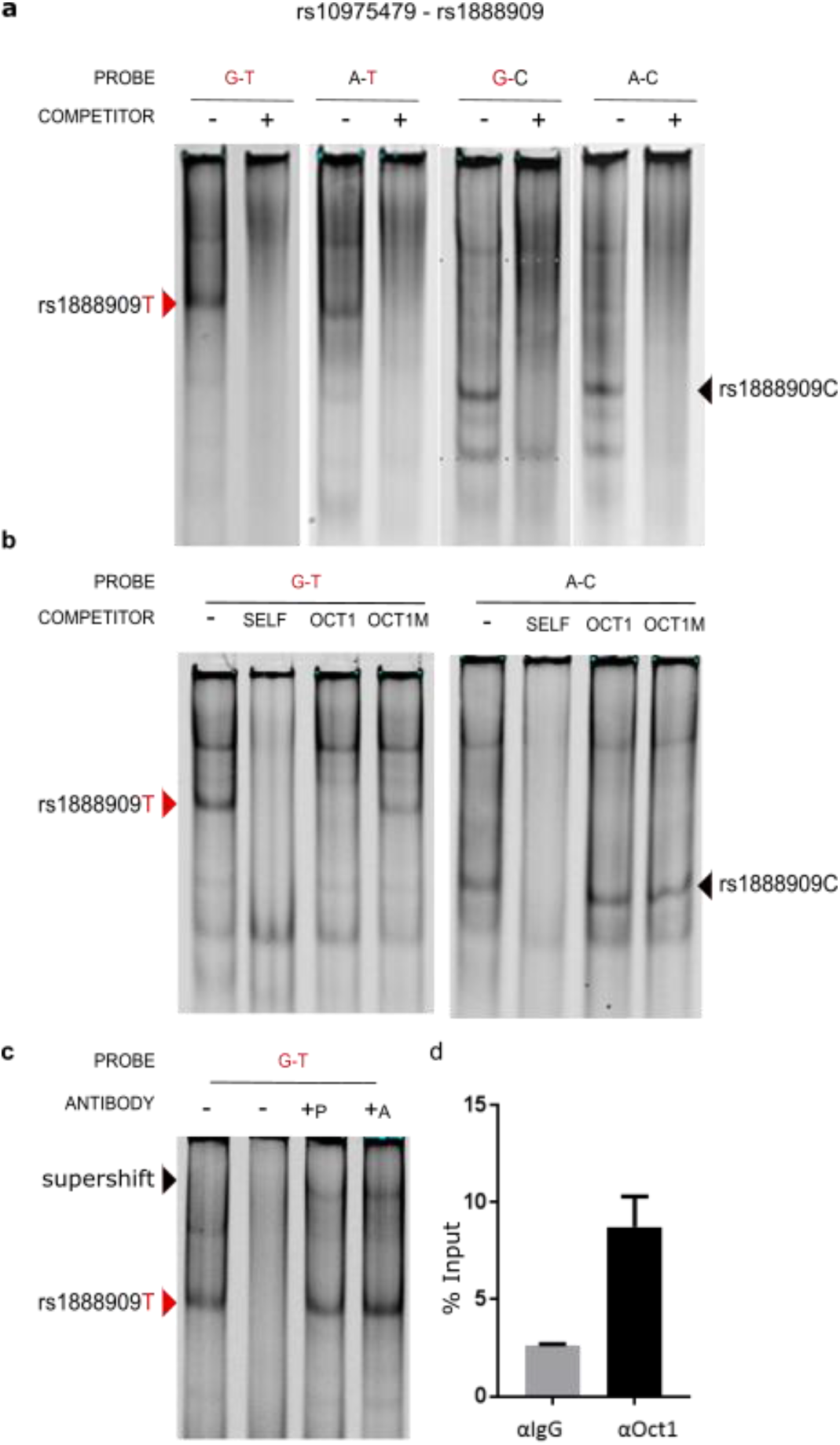
Regulatory region containing the risk allele rs1888909T selectively binds OCT-1. **a** Radiolabeled probes carrying the risk (in red) and/or non-risk sequences for SNPs rs10975479 and rs1888909 were incubated with nuclear extract obtained from K562 cells. Different complexes formed by rs1888909 are marked by red or black arrows. **b** Cold competition assay with OCT-1 consensus (OCT1) or mutated OCT-1 (Oct1M) oligonucleotides. EMSA probes and oligo competitor (100x molar excess) are noted above each gel. **c** Supershift complex formation with addition of anti-OCT-1 antibody as indicated by the red arrow. +P indicates addition of probe with nuclear extract, followed by incubation with antibody. +A indicates incubation of extract with antibody followed by addition of probe. d Chromatin immunoprecipitation of H292 chromatin with anti-OCT-1 antibody shows percent enrichment to input chromatin compared to control IgG antibody. Results are representative of 3 independent experiments

To identify nuclear proteins that bind to the rs1888909 (T) vs rs1888909 (C) probes, we isolated 3 bands from the non-risk rs1888909 (C) lane, and 2 bands from the risk rs1888909 (T) lane for mass spectrometry analysis (Supplementary Fig. 7). As a control, we isolated the same regions in the lane where the lysate had been incubated with a cold probe competitor. After filtering out the proteins that were also found on the control lane and the nuclear proteins that aren’t known to bind to DNA, we identified 3 transcription factors bound only to the non-risk probe, NFE2, TFCP2 and FOXL2 and 4 transcription factors bound only to the risk probe, OCT-1 (POU2F1), FOXP1, STAT3 and STAT5b. We then used the UniPROBE protein array database^31^ to select the transcription factors known to bind to at least one octamer containing a risk or non-risk allele. This analysis resulted in one transcription factor: OCT-1 (POU2F1), which bound to the EMSA probe and was differentially bound to the risk allele. OCT-1 was selected as our primary candidate for further investigation.

To confirm OCT-1-specific binding to the risk probe we first used a cold competition assay and demonstrated that the band specifically competed with an OCT-1 canonical DNA binding motif, but not with a mutated oligonucleotide (Fig. 5b, left panel). Conversely, binding to the non-risk A-C probe was not competed by the consensus or mutated OCT-1 oligonucleotides, demonstrating specificity for OCT-1 association with only the risk allele (Fig. 5b, right panel). Further, an OCT-1 specific antibody supershifted the nuclear complex formed with the rs1888909 (T) probe (Fig. 5c). We were able to visualize this shift independently of the order of incubation of the nuclear extract with probe or antibody, suggesting robust protein binding. We then performed chromatin immunoprecipitation (ChIP) followed by qPCR to demonstrate enrichment of OCT-1 binding to the DNA region containing the rs18889099 (T) (Fig. 5d). These experiments provide strong evidence that OCT-1 binds differentially to the risk allele and directs the formation of differential allelic nuclear complexes at the rs1888909 variant.

### Asthma-associated SNPs are associated with *IL33* mRNA and IL-33 protein levels

Finally, we tested the functional consequences of the GWAS SNPs rs1888909, rs10975479 and rs992969 on *IL33* mRNA and IL-33 protein abundance. The first two SNPs are within the 5 kb region. The latter, rs992969, is located outside of this region, but was previously reported to be associated with *IL33* expression in bronchial epithelial cells from primarily non-Hispanic white subjects^32^. SNP rs992969 is in high LD with rs1888909 in European ancestry populations (*r*^2^=1 in CEU), but less so in African American populations (*r*^2^=0.45 in ASW), and in low LD with rs10975479 in both populations (*r*^2^=0.56 and 0.13, respectively) (Supplementary Figure 1 and Supplementary Table 1). We used RNA-seq data from endobronchial brushings obtained from 123 asthmatic and non-asthmatic adult subjects, mostly of European ancestry (Fig. 6a), and from nasal epithelial cell brushings from 189 African American children from high risk asthma families (Fig. 6b). In both populations, carriers of one or two copies of the rs1888909 (T) risk allele had significantly higher *IL33* transcript levels compared to non-carriers of this allele. Genotypes at rs992969 were more modestly associated with transcript levels and only significant in the African American samples after correcting for multiple (3) tests; rs10975479 was not associated with *IL33* transcript abundance in either sample (Fig. 6a-b, Supplementary Table 3). These results are consistent with the EMSA data suggesting that allelic variants of rs1888909 result in differential protein binding. The stronger effects of rs1888909 on *IL33* expression in both populations suggest that rs1888909 is the causal variant in this region and that associations with other variants (such as rs992969) in GWAS and in gene expression studies were due to LD with the causal variant.

**Figure 6.**
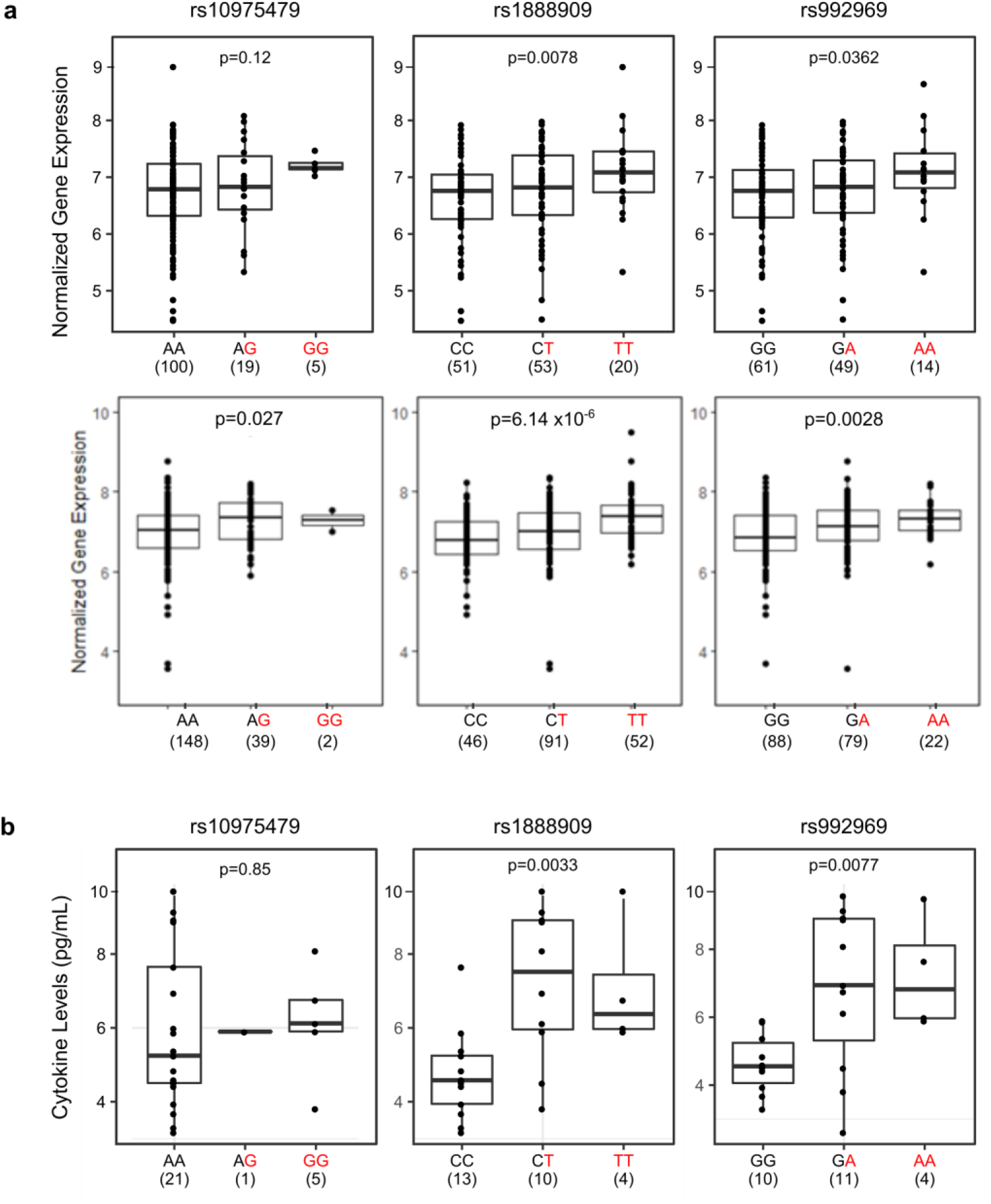
The rs1888909(T) and rs992969(A) alleles are associated with increased *IL33* expression and IL-33 protein levels. Comparison of *IL33* expression between genotypes for SNPs rs10975479, rs1888909 and rs992969 from **a** endobronchial brushings from 124 asthmatic and non-asthmatic adult subjects, mostly of European ancestry, and **b** nasal epithelial cells from 189 African American children from high risk asthma families. **c** Comparison of IL-33 cytokine levels between genotypes for SNPs rs10975479 (n=26), rs1888909 (n=27) and rs992969 (n=25) measured in plasma from Hutterite children (all European ancestry). The asthma-associated risk allele at each SNP is highlighted in red (x axis). The number of subjects per group is shown below the genotype. Boxes indicate the interquartile range, whiskers represent the 95% confidence intervals. Statistical significance was determined using an additive linear mode

We observed similar patterns of association in studies of IL-33 protein levels in plasma from 30 children^33^ of European ancestry (Fig. 6b). Children who carried the asthma risk alleles at rs1888909 or rs992969 had more IL-33 protein compared to children not carrying these alleles. There was no association between genotype at SNP rs10975479 and IL-33 protein levels.

The aggregate of our mouse transgenic assays, *in vivo* and *in vitro* reporter assays, and EMSA collectively supported a role for rs1888909 as a causal variant for the association with *IL33* expression. The associations between the risk alleles and increased *IL33* transcript levels and IL-33 protein abundance further support a role for this region in regulating the *IL33* gene and pointing to rs1888909 as the causal variant in this region, corroborating predictions from our *in vitro* and in vivo studies.

## Discussion

We described an integrated pipeline to fine-map and functionally annotate the asthma-associated locus that includes the *IL33 gene*. We used the LD structure at this locus across populations of different ethnicities combined with a Bayesian fine-mapping tool to define a critical 20 kb genomic interval containing candidate causal SNPs for the asthma association. Epigenetic signatures further reduced this region to 5 kb, which we demonstrated to have a crucial role in the development of chromatin loops creating contacts between *IL33* promoters and regulatory elements within the critical interval to control *IL33* expression. Associations between rs1888909 (T) allele copies and increased *IL33* mRNA and IL-33 protein levels further suggests that this regulatory variant mediates the development of asthma among individuals carrying this risk allele.

The 5 kb region that we delineated and characterized in this study defies the standard definition of regulatory elements. A loss of function assay showed that this region is necessary for *IL33* expression *in vivo*, suggestive of an enhancer function. However, we failed to detect enhancer activity in reporter assays in cells where the region displays chromatin markers associated with enhancers. Moreover, ENCODE profiled the binding of hundreds of transcription factors across multiple cell types and reports no binding of transcription activators, transcriptional co-factors, RNA PolII or other factors usually associated with enhancers within this 5 kb region. Rather, the region is bound by CTCF and cohesin in most cell lines assayed by ENCODE. We showed that this region possesses enhancer blocking activity *in vivo* and *in vitro*, reminiscent of insulator activities. Nevertheless, the region does not appear to behave as a classically described insulator, as we fail to see a change in chromatin topology and long-range interactions in the locus secondary to the deletion of the 5 kb region in the human BAC, which are typically seen when insulators are deleted ^30^. While we cannot ascribe a well-defined category for this regulatory element, we were able to dissect its functional properties, to fine-map a variant that is likely causal to the association with asthma and to identify the molecular effector binding to this regulatory variant.

CTCF plays a critical role in chromatin loop formation and participates in demarcation of Topological Association Domains (TADs). Loop formation within TADs facilitates contacts between specific genes and enhancers, allowing for appropriate temporal and tissue-specific expression^34^. We observed that the 5 kb risk haplotype carrying the asthma-associated allele possessed reduced enhancer blocking function in both *in vitro* and *in vivo* assays, and identified a smaller 1 kb fragment which maintained enhancer blocking activity. Interestingly, the loss of a 400bp region within this 1 kb fragment, harboring 2 asthma-associated SNPs, led to a significant loss of function, demonstrating a requirement for factors binding in that region for its proper regulatory activity.

We identified OCT-1 as a transcription factor that binds differentially to the risk allele of rs1888909, thus implicating OCT-1 in the function(s) of this 5 kb region. OCT-1 can regulate gene expression both positively and negatively. It has been shown to bind enhancers and regulatory regions upstream of multiple cytokines, including IL-3^35^, IL-12p40^36^, IL-13^37^, IL-4^38^ and IL-17^39^. The mechanisms by which OCT-1 functions are strikingly diverse, and there are numerous studies reporting the coordinated activity of CTCF and OCT-1. For example, at the *IL17* gene cluster on chromosome 3, OCT-1 and CTCF facilitate long-range associations with the IL-2 locus in naïve T cells. In a parallel fashion, it is possible that OCT-1 coordinates with CTCF to direct *IL33* expression at the locus through creation of appropriate chromatin interactions between enhancers and the *IL33* promoter. The distinct binding complexes formed on the risk and non-risk alleles at the rs1888909 genomic region may be responsible for alterations in these interactions, which in turn may lead to altered *IL33* expression levels.

While our data support a role of rs1888909 in the regulation of *IL33* expression and its primary role with asthma risk, our fine mapping results identified two other potential causative sets of SNPs associated with asthma at the *IL33* locus, upstream and downstream to the region that we studied (Fig1b). This suggests that other variants in this locus may have independent effects that alter the function of other regulatory elements and potentially controls expression of *IL33* or other genes. Future studies will be needed to further dissect the genetic architecture at this complex locus and address this hypothesis.

In summary, we identified a small 5 kb noncoding interval which is integral to *IL33* expression in several cell types. We propose a model in which CTCF mediates interaction of the 5 kb region to the *IL33* promoters through the formation of chromatin loops. The interplay between the distinct regulatory elements in the locus promotes spatial and temporal-specific regulation which is affected by OCT-1 binding to rs1888909. These results together with the increased *IL33* expression observed in humans with the risk allele offer a plausible mechanistic explanation for the association of these variants and asthma risk. A deeper characterization of these mechanisms will not only enhance our understanding of *IL33* regulation but may also open new strategies in directing future studies, as well as other common diseases in which dysregulation of IL-33 plays a role.

## Materials and Methods

### Experimental animals

Generation of hI*L33*^Crm^ BAC and h*IL3*3^Crm^ 5Kdel transgenic mice was performed by the University of Chicago Transgenic Core Facility. Modified DNA was diluted to a concentration of 2 ng/μl and used for pronuclear injections of CD1 embryos in accordance with standard protocols approved by the University of Chicago.

### Red/ET BAC modification

BAC RP11725F15, obtained from the BACPAC Resource Center (Oakland, CA), was modified in vitro using the RED/ET recombination kit (Gene Bridges, Heidelberg Germany) according to the manufacturer’s instructions.

Crimson-kanamycin cassette was PCR amplified from vector pE2-Crimson-N1 vector. The reporter cassette was inserted in frame to the ATG of the human *IL33* gene, to replace the second exon of the gene, while maintaining boundaries and flanking regions fully intact. Primers containing 50-bp BAC-homology arms used for generation of the recombination cassette were as follow: IL33_CrimsonKanF: TTGAGACAAATGAACTAATATTATATTTTAATCCAACAGAATACTGAAAAATGGATAGCACTGAGAACGTCA; IL33_CrimsonKanR, GCGTAAAACATTCAGAGATAACTTAAGTCCTTACTTCCCAGCTTGAAACATCAGAAGAACTCGTCAAGAAGG. Successful recombinants were screened for homologous insertion using the following external/internal primers IL33F-Crm5’, AGCCACAGTTGTTTCCGTTT; IL33R-Crm5’, TTGAGGTAGTCGGGGATGTC and IL-33F-Crm3’, ATCGCCTTCTATCGCCTTCT; IL33R, Crm3’, TGTGGAGCAAAAAGTGGTTG.

The asthma-associated 5 kb interval (chr9: 6,194,500-6,199,500; hg19) was deleted using RED/ET recombination kit (Gene Bridges). The 5 kb region of interest was replaced by the ampicillin gene using the following primers containing 50 pb homology arms flanking the BAC region to be deleted. IL33InsDEL_F, GCACACCTGTAAGTCTCTGCATTTTGCCACTTATACAACTTCATCTTTGAGTGGCACTTTTCGGGGAAATG; IL33InsDEL_R, AAACTTACATCAAATAAAATCTCAACACAGAATTCATACATGTCAACATACTCGAGGCTAGCTCTAGAAGTC. All correctly modified Bacs were verified by fingerprinting and sequencing. BAC DNA was extracted using the Nucleobond PC20 kit (Macherey-Nagel) and diluted for pronuclear injection. BAC copy number was determined as previously described ^40^

### Quantitative RT-PCR

RNA from flash frozen tissues were extracted from control and insulator deleted mice using TRI-reagent (Sigma). cDNA was generated using Super Script II (Thermo). IL33 primers (5’GCCAAGCTGCAAGTGACCAA 3’; 5’GCCTTGGAGCCGTAGAAGAA 3’) and HPRT housekeeping gene primes were used to quantify *IL33* mRNA expression levels. RNA samples with no reverse transcriptase added were used to test for genomic contamination.

### Tissue preparation, Immunofluorescence Staining, and Microscopy

Mouse lymph nodes were embedded into blocks with Optimal Cutting Temperature compound (OCT 4583) and stored at −80^°^C. Frozen tissues were sliced into sections 5μm thick and dried onto slides overnight. Sections were fixed, permeabilized, quenched and blocked. Tissues were immunostained with the primary antibodies rat anti-mouse CD31:Biotin (clone 390, Biolegend, San Diego, CA, USA) and Living Colors DsRed Polyclonal Antibody (rabbit anti-E2-Crimson, Clontech [now Takara Bio USA], Mountain View, CA, USA). Sections were washed and stained with the secondary antibodies Streptavidin:Alexa Fluor 488 (Biolegend, San Diego, CA, USA) and Goat anti-Rabbit IgG:Alexa Fluor 633 (Life Technologies, Thermo Fisher, Waltham, MA, USA), along with the nucleic acid stain Hoechst 33342 (Life Technologies, Thermo Fisher, Waltham, MA, USA). Coverslips were set with ProLong Diamond Antifade Mountant (Life Technologies, ThermoFisher, Waltham, MA, USA). Imaging was performed at the University of Chicago Integrated Light Microscopy Facility. Images were captured with a Leica TCS SP2 laser scanning confocal microscope (Leica Microsystems, Inc., Buffalo Grove, IL, USA) using a 63x/1.4 UV oil immersion objective and LAS_AF acquisition software (Leica Microsystems, Inc., Buffalo Grove, IL, USA). Further processing was completed using ImageJ software (National Institutes of Health, Bethesda, MD, USA).

### Promoter capture in situ Hi-C

Promoter capture Hi-C (PCHiC) data for human lymphoblast cells (LCL) was obtained from NCBI GEO (accession number: GSE79718), mapped to human genome assembly hg19, and processed as previously described^41, 42^. Interactions were visualized using then WashU Epigenome Browser.

### Circularized Chromosome Conformation Capture sequencing (4C-seq)

4C-seq assays were performed as previously reported^9^. 20 million bone marrow derived dendritic cells were obtained from control and insulator deleted mice. Immediately before 4C-seq library preparation, cells were cross-linked with 1% fresh formaldehyde for 15 minutes and treated with lysis buffer (10 mM Tris-HCl pH 8, 10 mM NaCl, 0.3% IGEPAL CA-630 (Sigma-Aldrich), 1X complete protease inhibitor cocktail (Roche). Nuclei were digested with Csp6-I endonuclease (Life Technologies, ThermoFisher, Waltham, MA) and ligated with T4 DNA ligase (Promega, Madison, WI) for 12 hours at 4°C, followed by reverse crosslinking at 65°C for 12 hours with proteinase K. Subsequently, NlaIII endonuclease (New England Biolabs) was used in a second round of digestion, and the DNA was ligated again. Specific primers were designed near the 5 kb region of interest. Viewpoint fragment ends (fragends) coordinates was chr9:6,186,164-6,186,468. Primers used for 4C-Seq experiments are as follow: 4C*mIL33IL33*Enh_nonread, 5’ CAAGCAGAAGACGGCATACGAACAACTTCACTCAGAGGCATG 3’ and 4C*mIL33IL33*Enh_Read 5’ AATGATACGGCGACCACCGAACACTCTTTCCCTACACGACGCTCTTCCGATCTGAGATGGCGCCACTGTAC 3’

### Constructs preparation

PCR fragments of genomic DNA from a risk and non-risk cell line were cloned into the pDONR vector (Life Technologies, ThermoFisher, Waltham, MA) and sequenced by the University of Chicago DNA Sequencing and Genotyping facility. DNA was subcloned into the enhancer barrier vector^28^ (gift from Dr. Laura Elnitski, NHGRI) or pGL4.23 for enhancer assays (Promega, Madison, WI). Constructs were prepared using the Plasmid MidiPrep Kit (Qiagen, Hilden, Germany) and re-sequenced to confirm genotype.

### Luciferase assay

K562 erythroleukemic were transfected using TransIT-2020 Transfection Reagent (Mirus Bio, Madison, WI). Briefly, 10^5^ cells per well were plated 24 hours prior to transfection. The transfection mix included 0.5μg of barrier or enhancer plasmid, 10ng hprl normalization control plasmid, and 1.5μl transfection reagent. TeloHaecs were transfected using JetPRIME transfection reagent (Polyplus). 20,000 cells per well (24 well plate) were plated and transfect with 250ng of test plasmid DNA and 25 ng of hprl normalization control plasmid. After 48 hours cells were harvested and assayed for luciferase with a 20/20^n^ luminometer (Turner Biosystems, now Fisher Scientific, Hampton, NH) using the Dual Luciferase Reporter Assay System (Promega, Madison, WI). Firefly luciferase was normalized to the Renilla luciferase. A minimum of three independent transfections using 3 different DNA preparations were assayed.

### Zebrafish transgenesis

The Tol2 vector contains a strong midbrain enhancer, a Gateway entry site and the cardiac actin promoter controlling the expression of EGFP, and was developed to screen for insulator activity^29^. Each candidate sequence was recombined between the midbrain enhancer and the cardiac actin promoter. As a reference, the empty backbone was used (INS-zero). One cell-stage embryos were injected with 3–5 nL of a solution containing 25 nM of each construct plus 25 nM of Tol2 mRNA. Embryos where then incubated at 28°C and EGFP expression was evaluated 24 hpf. The midbrain/somites EGFP intensity ratio was quantified using ImageJ freeware and was directly proportional to the enhancer-blocking capacity. As a positive control, the chicken beta-globin insulator 5HS4 was used. Each experiment was repeated independently and double-blinded to the operators.

### EMSA

Nuclear extracts were prepared from K562 cells using NE-PER Nuclear and Cytoplasmic Extraction Kit (Thermo Fisher, Waltham, MA) supplemented with HALT Protease Inhibitor Cocktail and PMSF (Thermo Fisher, Waltham, MA). Protein concentrations were determined with the BCA kit (Thermo Fisher, Waltham, MA). Oligonucleotides to be used as probes were synthesized with a 5’IRDye 700 modification (IDT). Binding reactions were performed using the Odyssey EMSA kit (LI-COR Biosciences, Lincoln, NE) and contained 5μg nuclear extract, 2.5nM labeled probe, and when noted, 100x excess unlabeled oligonucleotide. For supershift assays, 2μg Oct-1 antibody (Santa Cruz Biotechnology, Dallas, TX, sc-232) was added and incubated for another 20 minutes. Reaction mixtures were run on a 4% nondenaturing polyacrylamide gel and analyzed with the LI-COR Odyssey Imaging System.

### Oligonucleotide/Probe sequences

479G:909 TCTGATGCAGAACA<u>G</u>CAATGTGTTTTCCA<u>T</u>GTGCACTTGGTC

479G:909 CCTGATGCAGAACA<u>G</u>CAATGTGTTTTCCA<u>C</u>GTGCACTTGGTC

479A:909 TCTGATGCAGAACA<u>A</u>CAATGTGTTTTCCA<u>T</u>GTGCACTTGGTC

479A:909 CCTGATGCAGAACAACAATGTGTTTTCCACGTGCACTTGGTC

Consensus Oct1 TGTCGAATGCAAATCACTAGAA

Mutant Oct1 TGTCGAATGCAAGCCACTAGAA.

### Mass Spectrophotometry and protein identification

The gel sections to be analyzed from the EMSA gel were excised, washed and destained using 100mM NH4HCO3 in 50% acetonitrile (ACN). Sections then underwent reduction, alkylation, and trypsinization. Peptides were extracted with 5% formic acid, followed by 75% ACN in formic acid, and cleaned up with C-18 spin columns (Pierce). The samples were analyzed via electrospray tandem spectrometry (LC-MS/MS) on a Thermo Q-Exactive Orbitrap mass spectrometer at the Proteomics Core at Mayo Clinic, Rochester, Minnesota. Tandem mass spectra were extracted and then analyzed by Mascot and X!Tandem algorithms. Scaffold v4.8.4 (Proteome Software Inc., Portland,OR) was used to validate the protein identifications. Peptide identifications were accepted if they could be established at greater than 98% probability to achieve an FDR less than 1.0%.

### Candidate Transcription factor analysis

Panther pathway analysis database^43^ was used to filter for DNA-binding proteins. Jaspar (http://jaspar.genereg.net) and Alggen-promo (http://alggen.lsi.upc.es) databases were used to screen these factors for known DNA binding sites. We downloaded binding data for random octamers from UniProbe (http://thebrain.bwh.harvard.edu/pbms/webworks_pub_dev/downloads.php) and identified all transcription factors that bound at least one octamer containing a risk or non-risk allele.

### Chromatin Immunoprecipitation

ChIP assays were performed using the Millipore ChIP assay kit and protocol. 2×10^6^ H292 cells (heterozygous for the risk allele) were fixed with 1% formaldehyde. Chromatin was sonicated using a diagenode BioRuptor. Lysates were incubated overnight with 5ug of Oct-1 (Santa Cruz, sc-232x) or an IgG control (Santa Cruz, sc-2027). Following the washes, elution of chromatin complexes and reversal of crosslinks, DNA was recovered using Qiagen pcr purification kit. Input DNA was also processed. QPCR was performed using the primers: F: 5’GCCTCTGGTCTCAGTGGATA3’ and R: 5’CTGCTCATAGGAGACACAGTAAAG3’.

### Gene expression and genotype studies

RNA-seq data from airway epithelial cells were available from two sources. The first was from bronchial epithelial cells sampled from adults (44 European American, 48 African American), with and without asthma (76 cases, 60 controls), who participated in a study of asthma in Chicago. Procedures for bronchoscopy, cell and RNA processing, and genotyping have been previously described^41^. For this study, normalized gene expression counts were adjusted for age, sex, current smoking status, sequencing pool, the first three ancestry PCs. Linear regression considering additive genotype effects on gene expression was performed using limma in R (v3.3.3). *P*-values were adjusted for 3 tests; *P*<0.016 was considered significant. All subjects provided written informed consent; these studies were approved by University Chicago Institutional Review Board.

The second source was nasal epithelial cells sampled from 246 African American children (125 cases, 121 controls) from a birth cohort of children at high risk for asthma^44^. Procedures for nasal brushing and cell processing followed standard procedures^45^. Genotypes were determined using Illumina Multi-Ethnic Genotyping Array (MEGA), and processed using standard QC^41^. To test for association between genotypes for three SNPs and *IL33* transcript levels, we used an additive effects linear model, including as covariates sex, study site, batch id, epithelial cell proportion and 12 latent factors in the epithelial cell studies. Latent factors were included to correct for unwanted variation^46^. *P*-values were adjusted for 3 tests; *P*<0.016 was considered significant. Parents of all children provided written informed consent, and children provided written assent, for genetic studies; these studies were approved by University Chicago Institutional Review Board.

### IL-33 cytokine and genotype studies

Blood for cytokine studies was drawn from 30 Hutterite children during trips to South Dakota, as previously described^47^. Written consent was obtained from the parents and written assent was obtained from the children. The study was approved by the institutional review boards at the University of Chicago. Briefly, The Milliplex Map Human TH17 Magnetic Bead Panel (Millipore, Burlington, MA) was used to measure IL-33 levels in thawed supernatants at the University of Chicago Immunology Core facility using standard protocols, as previously described.^48^

Associations between genotypes at rs1888909, rs992969 and rs10975479 and normalized IL-33 cytokine abundance were tested using a linear model, assuming additive effects. Prior to analysis, voom-transformed gene expression counts were adjusted for age, sex, current smoking status, sequencing pool and the first three ancestry PCs using the function removeBatchEffect from the R package limma^49^. Linear regression between the genotypes and IL-33 levels was performed with the FastQTL software package^50^. One subject had poor IL-33 cytokine data and was removed. Of the remaining subjects, 27 subjects for rs10975479 and rs1888909 and 25 subjects for rs992969 had good quality genotype calls and were retained. Linear regression between the genotypes and IL-33 cytokine levels was performed using limma, adjusting for sex and age. Genotypes were obtained using PRIMAL^51^, an in-house pedigree-based imputation tool that imputes variants from whole genome sequences from 98 Hutterite individuals to >1600 individuals who were genotyped with an Affymetrix genotyping array.

## Acknowledgements

We thank Don Wolfgeher, from the University of Chicago’s Proteomics Core Facility, for assistance with sample preparation and analysis of the mass spectrophotemetry; Dr. Laura Elnitski (NHGRI/NIH, Bethesda,MD), for generously supplying the enhancer barrier plasmids. Dr. Benjamin Glick (University of Chicago) for supplying the E2-Crimson reporter vector.

## REFERENCES

1. Polderman, T.J. et al. Meta-analysis of the heritability of human traits based on fifty years of twin studies. Nature genetics 47, 702–709 (2015).

2. Pividori, M., Schoettler, N., Nicolae, D.L., Ober, C. & Im, H.K. Shared and distinct genetic risk factors for childhood-onset and adult-onset asthma: genome-wide and transcriptome-wide studies. Lancet Respir Med 7, 509–522 (2019).

3. Schmitz, J. et al. IL-33, an interleukin-1-like cytokine that signals via the IL-1 receptor-related protein ST2 and induces T helper type 2-associated cytokines. Immunity 23, 479–490 (2005).

4. Prefontaine, D. et al. Increased expression of IL-33 in severe asthma: evidence of expression by airway smooth muscle cells. J Immunol 183, 5094–5103 (2009).

5. Kearley, J., Buckland, K.F., Mathie, S.A. & Lloyd, C.M. Resolution of allergic inflammation and airway hyperreactivity is dependent upon disruption of the T1/ST2-IL-33 pathway. Am J Respir Crit Care Med 179, 772–781 (2009).

6. Savinko, T. et al. IL-33 and ST2 in atopic dermatitis: expression profiles and modulation by triggering factors. J Invest Dermatol 132, 1392–1400 (2012).

7. Kamekura, R. et al. The role of IL-33 and its receptor ST2 in human nasal epithelium with allergic rhinitis. Clin Exp Allergy 42, 218–228 (2012).

8. Kottyan, L.C. et al. Genome-wide association analysis of eosinophilic esophagitis provides insight into the tissue specificity of this allergic disease. Nature genetics 46, 895–900 (2014).

9. Smemo, S. et al. Obesity-associated variants within FTO form long-range functional connections with IRX3. Nature 507, 371–375 (2014).

10. van den Boogaard, M. et al. A common genetic variant within SCN10A modulates cardiac SCN5A expression. J Clin Invest 124, 1844–1852 (2014).

11. Rusu, V. et al. Type 2 Diabetes Variants Disrupt Function of SLC16A11 through Two Distinct Mechanisms. Cell 170, 199–212 e120 (2017).

12. Small, K.S. et al. Regulatory variants at KLF14 influence type 2 diabetes risk via a female-specific effect on adipocyte size and body composition. Nature genetics 50, 572–580 (2018).

13. Wasserman, N.F., Aneas, I. & Nobrega, M.A. An 8q24 gene desert variant associated with prostate cancer risk confers differential in vivo activity to a MYC enhancer. Genome Res 20, 1191–1197 (2010).

14. Gallagher, P.G. et al. Mutation of a barrier insulator in the human ankyrin-1 gene is associated with hereditary spherocytosis. J Clin Invest 120, 4453–4465 (2010).

15. Moffatt, M.F. et al. A large-scale, consortium-based genomewide association study of asthma. N Engl J Med 363, 1211–1221 (2010).

16. Torgerson, D.G. et al. Meta-analysis of genome-wide association studies of asthma in ethnically diverse North American populations. Nature genetics 43, 887–892 (2011).

17. Bonnelykke, K. et al. A genome-wide association study identifies CDHR3 as a susceptibility locus for early childhood asthma with severe exacerbations. Nature genetics 46, 51–55 (2014).

18. Demenais, F. et al. Multiancestry association study identifies new asthma risk loci that colocalize with immune-cell enhancer marks. Nat Genet 50, 42–53 (2018).

19. Daya, M. et al. Association study in African-admixed populations across the Americas recapitulates asthma risk loci in non-African populations. Nature communications 10, 880 (2019).

20. Ferreira, M.A.R. et al. Genetic Architectures of Childhood-and Adult-Onset Asthma Are Partly Distinct. Am J Hum Genet 104, 665–684 (2019).

21. Zhu, X. & Stephens, M. Bayesian Large-Scale Multiple Regression with Summary Statistics from Genome-Wide Association Studies. Ann Appl Stat 11, 1561–1592 (2017).

22. Bycroft, C. et al. The UK Biobank resource with deep phenotyping and genomic data. Nature 562, 203–209 (2018).

23. Roadmap Epigenomics, C. et al. Integrative analysis of 111 reference human epigenomes. Nature 518, 317–330 (2015).

24. Consortium, E.P. A user's guide to the encyclopedia of DNA elements (ENCODE). PLoS Biol 9, e1001046 (2011).

25. Kim, S., Yu, N.K. & Kaang, B.K. CTCF as a multifunctional protein in genome regulation and gene expression. Exp Mol Med 47, e166 (2015).

26. Chen, D. & Lei, E.P. Function and regulation of chromatin insulators in dynamic genome organization. Curr Opin Cell Biol 58, 61–68 (2019).

27. Pichery, M. et al. Endogenous IL-33 Is Highly Expressed in Mouse Epithelial Barrier Tissues, Lymphoid Organs, Brain, Embryos, and Inflamed Tissues: In Situ Analysis Using a Novel Il-33-LacZ Gene Trap Reporter Strain. J Immunol 188, 3488–3495 (2012).

28. Petrykowska, H.M., Vockley, C.M. & Elnitski, L. Detection and characterization of silencers and enhancer-blockers in the greater CFTR locus. Genome Res 18, 1238–1246 (2008).

29. Bessa, J. et al. Zebrafish enhancer detection (ZED) vector: a new tool to facilitate transgenesis and the functional analysis of cis-regulatory regions in zebrafish. Dev Dyn 238, 2409–2417 (2009).

30. Flavahan, W.A. et al. Altered chromosomal topology drives oncogenic programs in SDH-deficient GISTs. Nature 575, 229–233 (2019).

31. Hume, M.A., Barrera, L.A., Gisselbrecht, S.S. & Bulyk, M.L. UniPROBE, update 2015: new tools and content for the online database of protein-binding microarray data on protein-DNA interactions. Nucleic Acids Res 43, D117–122 (2015).

32. Li, X. et al. eQTL of bronchial epithelial cells and bronchial alveolar lavage deciphers GWAS-identified asthma genes. Allergy (2015).

33. Stein, M.M. et al. Innate Immunity and Asthma Risk in Amish and Hutterite Farm Children. N Engl J Med 375, 411–421 (2016).

34. West, A.G., Gaszner, M. & Felsenfeld, G. Insulators: many functions, many mechanisms. Genes Dev 16, 271–288 (2002).

35. Duncliffe, K.N., Bert, A.G., Vadas, M.A. & Cockerill, P.N. A T cell-specific enhancer in the interleukin-3 locus is activated cooperatively by Oct and NFAT elements within a DNase I-hypersensitive site. Immunity 6, 175–185 (1997).

36. Zhou, L. et al. An inducible enhancer required for Il12b promoter activity in an insulated chromatin environment. Mol Cell Biol 27, 2698–2712 (2007).

37. Kiesler, P., Shakya, A., Tantin, D. & Vercelli, D. An allergy-associated polymorphism in a novel regulatory element enhances IL13 expression. Hum Mol Genet 18, 4513–4520 (2009).

38. Kim, K., Kim, N. & Lee, G.R. Transcription Factors Oct-1 and GATA-3 Cooperatively Regulate Th2 Cytokine Gene Expression via the RHS5 within the Th2 Locus Control Region. PLoS One 11, e0148576 (2016).

39. Kim, L.K. et al. Oct-1 regulates IL-17 expression by directing interchromosomal associations in conjunction with CTCF in T cells. Mol Cell 54, 56–66 (2014).

40. Chandler, K.J. et al. Relevance of BAC transgene copy number in mice: transgene copy number variation across multiple transgenic lines and correlations with transgene integrity and expression. Mamm Genome 18, 693–708 (2007).

41. Nicodemus-Johnson, J. et al. DNA methylation in lung cells is associated with asthma endotypes and genetic risk. JCI Insight 1, e90151 (2016).

42. Nicodemus-Johnson, J. et al. Genome-Wide Methylation Study Identifies an IL-13-induced Epigenetic Signature in Asthmatic Airways. Am J Respir Crit Care Med 193, 376–385 (2016).

43. Mi, H., Muruganujan, A., Ebert, D., Huang, X. & Thomas, P.D. PANTHER version 14: more genomes, a new PANTHER GO-slim and improvements in enrichment analysis tools. Nucleic Acids Res 47, D419–D426 (2019).

44. Gern, J.E. et al. The Urban Environment and Childhood Asthma (URECA) birth cohort study: design, methods, and study population. BMC Pulm Med 9, 17 (2009).

45. Poole, A. et al. Dissecting childhood asthma with nasal transcriptomics distinguishes subphenotypes of disease. J Allergy Clin Immunol 133, 670–678 e612 (2014).

46. McKennan, C. & Nicolae, D.L. Accounting for unobserved covariates with varying degree of estimability in high dimensional experimental data. Biometrika 106, 823–840 (2019).

47. Stein, M.M. et al. Innate Immunity and Asthma Risk in Amish and Hutterite Farm Children. N Engl J Med 375, 411–421 (2016).

48. Stein, M.M., Hrusch, C.L., Sperling, A.I. & Ober, C. Effects of an FcgammaRIIA polymorphism on leukocyte gene expression and cytokine responses to anti-CD3 and anti-CD28 antibodies. Genes Immun (2018).

49. Ritchie, M.E. et al. limma powers differential expression analyses for RNA-sequencing and microarray studies. Nucleic Acids Res 43, e47 (2015).

50. Ongen, H., Buil, A., Brown, A.A., Dermitzakis, E.T. & Delaneau, O. Fast and efficient QTL mapper for thousands of molecular phenotypes. Bioinformatics 32, 1479–1485 (2016).

51. Livne, O.E. et al. PRIMAL: Fast and Accurate Pedigree-based Imputation from Sequence Data in a Founder Population. PLoS Comput Biol 11, e1004139 (2015).

