## Supplementary Information for "Asthma-associated variants induce *IL33* differential expression through a novel regulatory region"

Aneas et al.

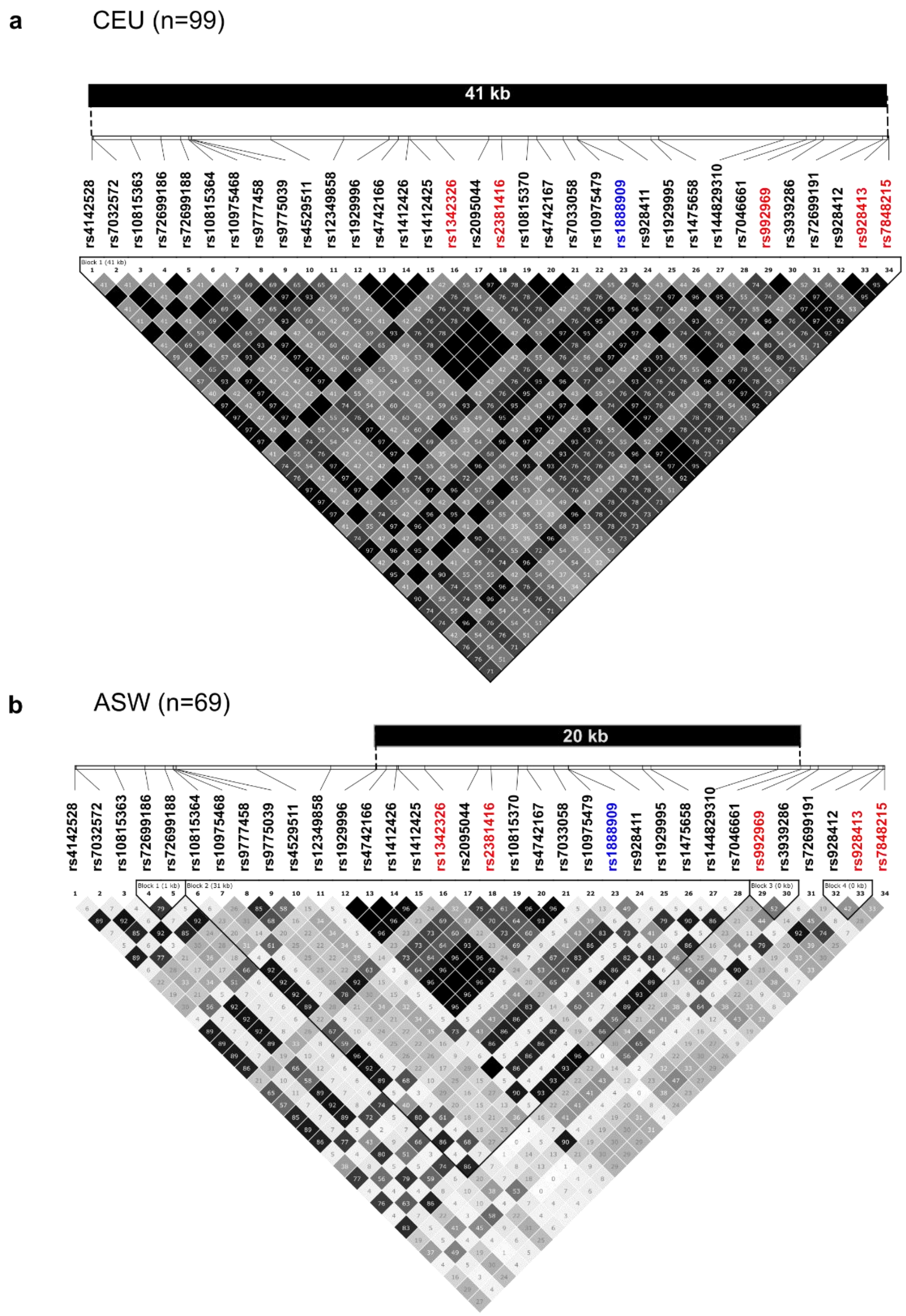

**Supplementary Figure 1**. 41kb LD blocks created with Haploview software using a MAF=0.1. Genomic coordinates are chr9: 6,172,380-6,213,488 (hg19). Position of the five lead SNPs (in red) from 7 GWAS studies are marked on the Caucasian (CEU) plot (**a**) and Americans with African ancestry from SW USA (ASW) plots **(b)**. The lead SNP rs1888909 in African Ancestry is shown in blue. The 34 SNPs showed in the plot are either in LD with one of five lead SNPs (in red) in European (r2≥8), African ancestry (r2≥0.4) or both populations.

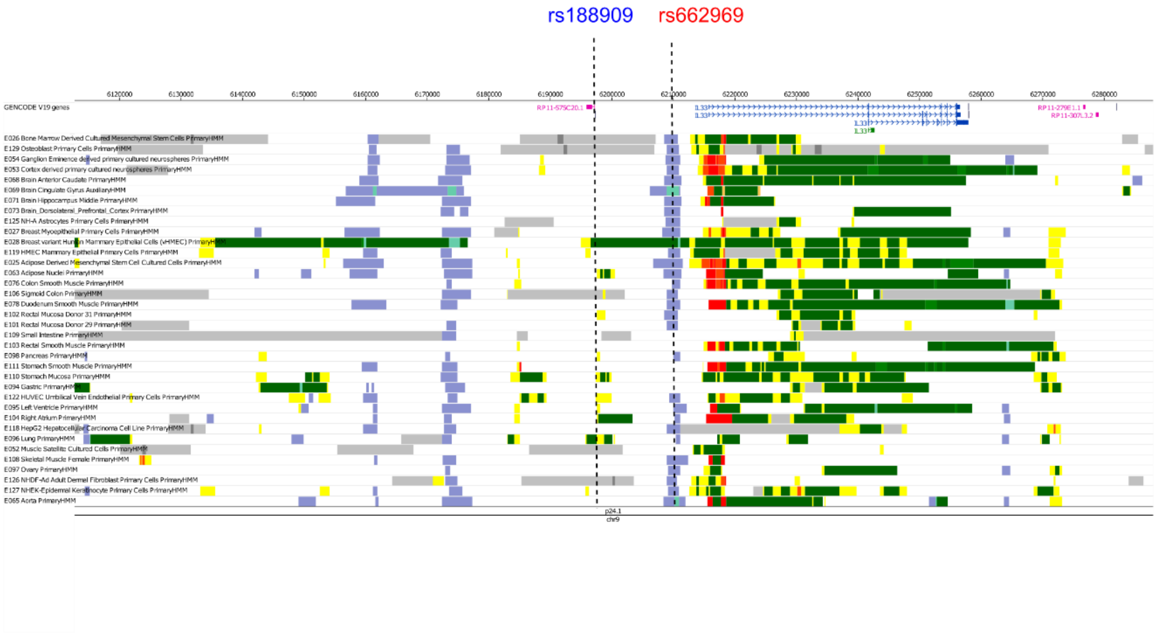

**Supplementary Figure 2. Roadmap Epigenome analysis at the *IL33* locus in 36 tissues.** rs662969 (in red) was the lead SNP in the combined multi- ancestry GWAS studies, while rs1888909 (in blue) was the lead in the African American/African Caribbean GWAS (marked in blue).

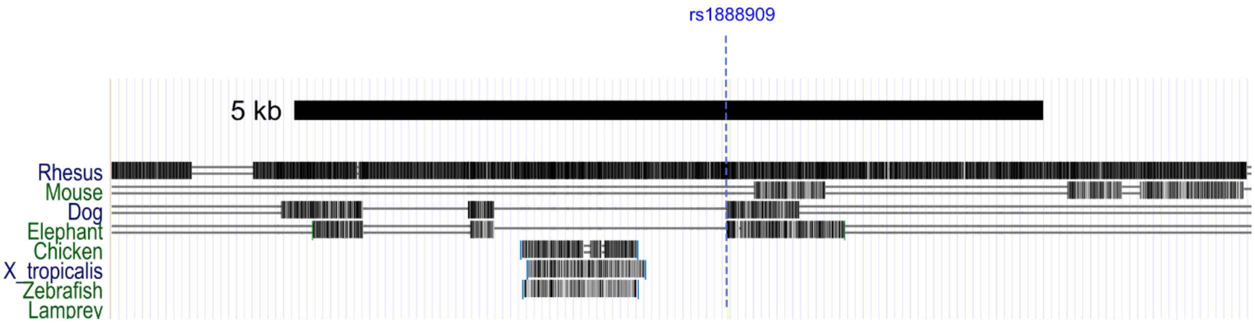

**Supplementary Figure 3. Multispecies conservation.** UCSC genome browser track (chr9:6,193,281-6,200,888, hg19) showing that the 5kb region is not conserved across species. A single line means no bases in the alignment, and double lines indicate one or more unaligned bases.

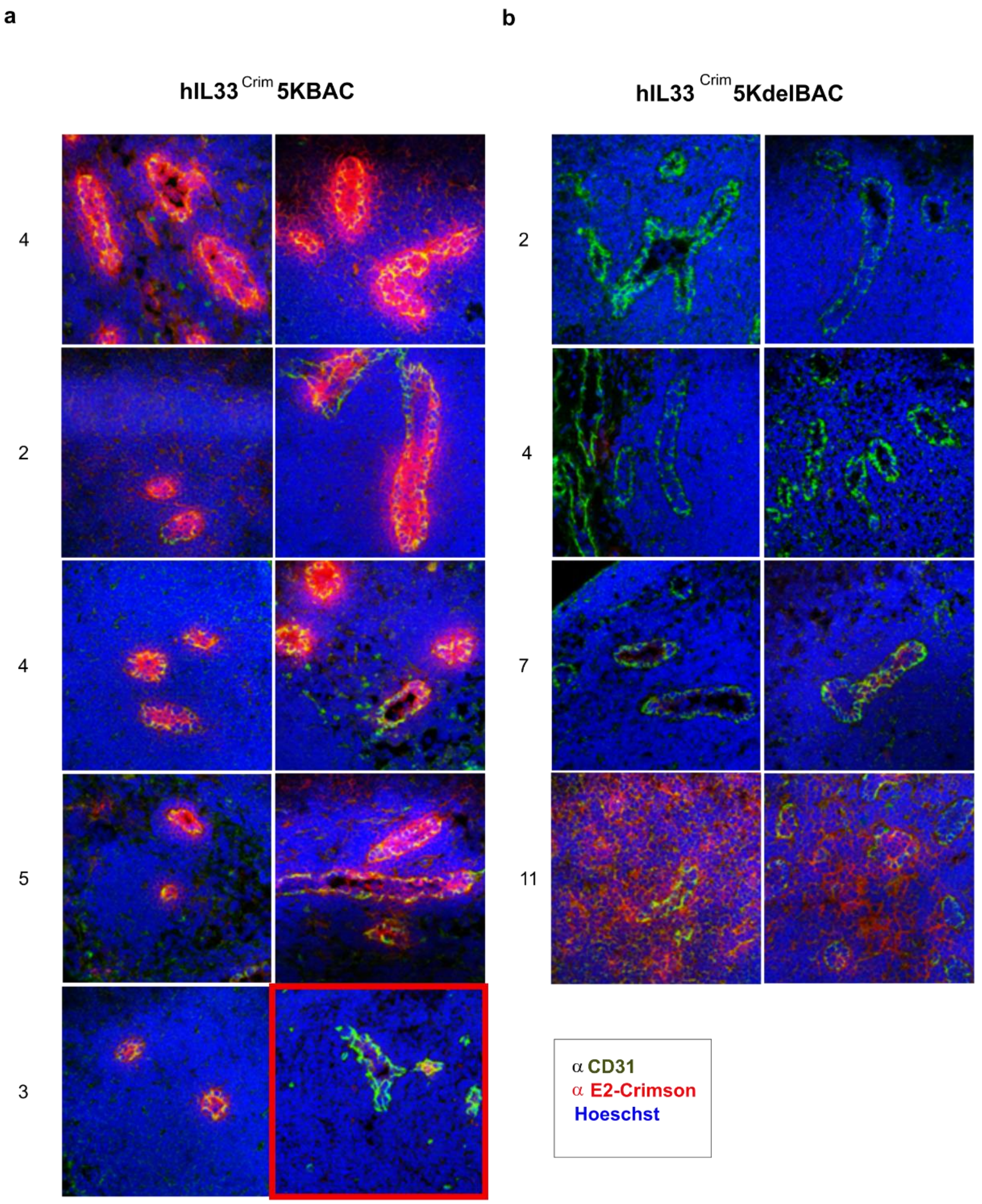

**Supplementary Figure 4. Immunohistochemistry of 5 h*IL33*$^{Crim}$ BAC transgenic founders** (**a**) and 4 h*IL33*$^{Crim}$BAC5kdel founders (**b**). Estimated BAC copy number is listed adjacent to each founder line. Lymph node sections from two mice for each line are shown. Red indicates Crimson expression. All sections except for one (red box) were stained with αE2-Crimson antibody and illustrates baseline Crimson expression. Green indicates staining with αCD31. Nucleic acid stain Hoescht is blue.

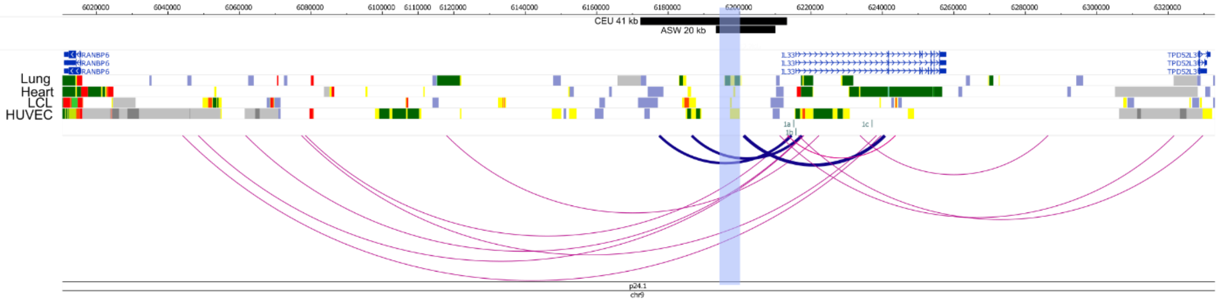

**Supplementary Figure 5. Promoter capture Hi-C (pcHi-C) in Lymphoblastoid cells (LCL).** Roadmap Epigenome data from heart (E095 Left ventricle primary HMM), lung (E096 Lung primary HMM), LCL (E116 GM128781 Lymphoblastoid cell primary HMM) and endothelial cells (E122 HUVEC Umbilical Vein Endothelial Primary Cells Primary HM). Chromatin state assignments from Roadmap Epigenome are indicated as: transcribed, green; active enhancer, yellow; active promoter, red; repressed, gray; heterochromatin, blue. Black bars correspond to the LD block upstream of exon 1 of the *IL33* gene. Blue shaded area correspond to the asthma-associated 5 kb region. Arcs depict interactions between *IL33* promoters and regions throughout the locus. Blue arcs depict interaction between the asthma-associated LD block region and both *IL33* promoters.

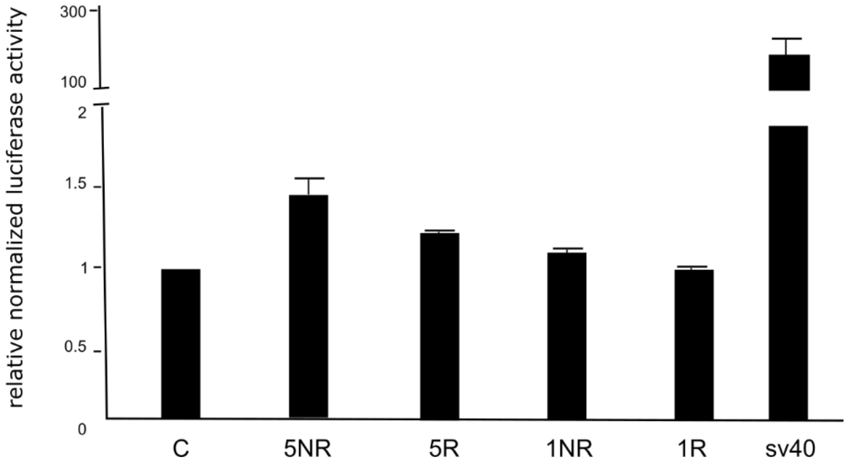

**Supplementary Figure 6. Luciferase enhancer assay.** Human aortic endothelial cells (TeloHaec) were transfected with a 5 kb (chr9: 6,194,500-6,199,500; hg19) or 1 kb (chr9: 6,197,000- 6,197,917; hg19) construct containing the non-risk (NR) or the risk (R) asthma- associated alleles. PGL3 control vector (Sv40) was used as positive control. The results are expressed as fold change between the test construct and the control vector normalized luciferase activity. Firefly luciferase/renilla luciferase provide the normalized luciferase activity for each vector.

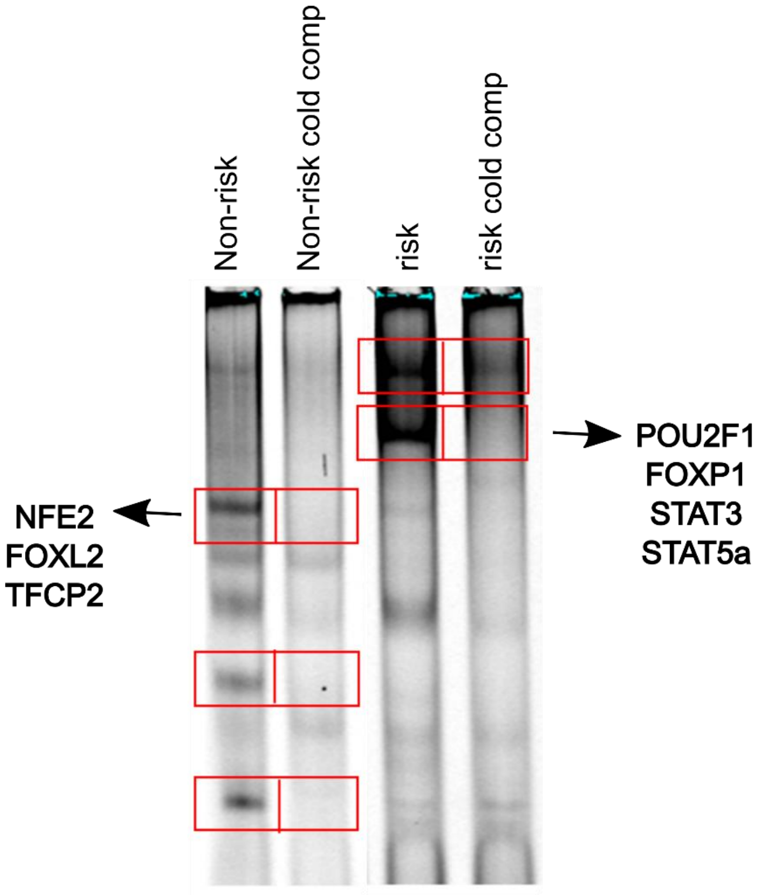

**Supplementary Figure 7**. EMSA assay showing outline of bands extracted for mass spectrophotometry. Unique transcription factors identified from the experiment are boxed. Note no transcription factors were identified in 2 bands from non-risk and 1 band from risk.

**Supplementary table 1**. SNPs in LD ($r^2$ ≥0.80) with at least one of the five lead SNPs (marked in red). A total of 26 SNPs spans a 41kb interval in the CEU population. SNPs in LD ($r^2$ ≥0.44) with the tag SNP rs1888909 in the African ancestry population (marked in blue) defined an LD block of 20 kb (chr9: 6,188,124-6,209,099) marked by the orange shaded area.

| chr | pos (hg19) | variant | Ref (risk) | Alt | GENCODE genes |
|---|---|---|---|---|---|
| 9 | 6172380 | rs7032572 | A | G | 43kb 5' of IL33 |
| 9 | 6175855 | rs72699186 | A | T | 40kb 5' of IL33 |
| 9 | 6176871 | rs72699188 | C | G | 39kb 5' of IL33 |
| 9 | 6177302 | rs10975468 | C | T | 39kb 5' of IL33 |
| 9 | 6177453 | rs9775039 | G | A | 38kb 5' of IL33 |
| 9 | 6178525 | rs144511606 | G | A | 37kb 5' of IL33 |
| 9 | 6185295 | rs12349858 | C | T | 31kb 5' of IL33 |
| 9 | 6187636 | rs1929996 | C | G | 28kb 5' of IL33 |
| 9 | 6188124 | rs4742166 | G | C | 28kb 5' of IL33 |
| 9 | 6188652 | rs1412426 | A | C,T | 27kb 5' of IL33 |
| 9 | 6190076 | rs1342326 | A | C | 26kb 5' of IL33 |
| 9 | 6192796 | rs2095044 | T | C | 23kb 5' of IL33 |
| 9 | 6193455 | rs2381416 | C | A | 22kb 5' of IL33 |
| 9 | 6194831 | rs10815370 | C | A | 21kb 5' of IL33 |
| 9 | 6195285 | rs4742167 | C | T | 21kb 5' of IL33 |
| 9 | 6196645 | rs7033058 | T | C | 19kb 5' of IL33 |
| 9 | 6197377 | rs10975479 | A | G | 18kb 5' of IL33 |
| 9 | 6197392 | rs1888909 | T | C | 18kb 5' of IL33 |
| 9 | 6201163 | rs1929995 | T | C | 15kb 5' of IL33 |
| 9 | 6208030 | rs144829310 | G | T | 7.8kb 5' of IL33 |
| 9 | 6209697 | rs992969 | A | G | 6.1kb 5' of IL33 |
| 9 | 6210099 | rs3939286 | T | C | 5.7kb 5' of IL33 |
| 9 | 6211813 | rs72699191 | T | C | 4kb 5' of IL33 |
| 9 | 6213148 | rs928412 | A | G | 2.7kb 5' of IL33 |
| 9 | 6213387 | rs928413 | G | A | 2.4kb 5' of IL33 |
| 9 | 6213468 | rs7848215 | C | T | 2.3kb 5' of IL33 |

**Supplementary table 2.** Sum of Single Effects (SuSiE)[22] to fine-map the associated region and identify credible sets (CS) of variants at the *IL33* locus with high probabilities of being causal. CS1 (n=6), CS2 (n=25) and CS3(n=28) are the 3 CS regions identified. SNPs marked in red were assigned the highest PIPs.

| CS | rsid | pos | pval | PIP |
|---|---|---|---|---|
| CS1 | rs992969 | 6209697 | 6.76E-42 | 0.391063 |
| CS1 | rs1888909 | 6197392 | 2.07E-41 | 0.208944 |
| CS1 | rs3939286 | 6210099 | 1.86E-41 | 0.204711 |
| CS1 | rs7848215 | 6213468 | 7.80E-42 | 0.085221 |
| CS1 | rs928412 | 6213148 | 4.38E-41 | 0.049622 |
| CS1 | rs2381416 | 6193455 | 1.25E-40 | 0.030186 |
| CS2 | rs1330124 | 6053098 | 2.36E-26 | 0.093386 |
| CS2 | rs343495 | 6069445 | 2.07E-26 | 0.088878 |
| CS2 | rs343487 | 6062269 | 4.54E-26 | 0.053872 |
| CS2 | rs343491 | 6064640 | 5.43E-26 | 0.047065 |
| CS2 | rs397505 | 6076823 | 1.99E-26 | 0.043794 |
| CS2 | rs343490 | 6064575 | 6.01E-26 | 0.043789 |
| CS2 | rs378952 | 6078146 | 1.91E-26 | 0.043609 |
| CS2 | rs10975413 | 6049843 | 6.49E-26 | 0.041792 |
| CS2 | rs343489 | 6064299 | 6.73E-26 | 0.039977 |
| CS2 | rs451974 | 6076844 | 2.40E-26 | 0.038493 |
| CS2 | rs340934 | 6081804 | 5.25E-26 | 0.037143 |
| CS2 | rs371454 | 6078614 | 3.08E-26 | 0.032018 |
| CS2 | rs343479 | 6054314 | 1.37E-25 | 0.028717 |
| CS2 | rs189348 | 6073194 | 5.39E-26 | 0.028704 |
| CS2 | rs380568 | 6055531 | 1.31E-25 | 0.028074 |
| CS2 | rs401834 | 6078991 | 3.68E-26 | 0.027771 |
| CS2 | rs343475 | 6073013 | 6.15E-26 | 0.026056 |
| CS2 | rs393556 | 6056468 | 1.59E-25 | 0.024546 |
| CS2 | rs10975418 | 6057011 | 1.73E-25 | 0.02344 |
| CS2 | rs340935 | 6080998 | 1.25E-25 | 0.022409 |
| CS2 | rs10975412 | 6049547 | 2.51E-25 | 0.021368 |
| CS2 | rs343496 | 6068077 | 1.08E-25 | 0.02007 |
| CS2 | rs343499 | 6059157 | 2.37E-25 | 0.019209 |
| CS2 | rs343476 | 6072597 | 1.18E-25 | 0.016354 |
| CS3 | rs10815391 | 6240235 | 5.67E-30 | 0.074113 |
| CS3 | rs10815392 | 6240236 | 5.67E-30 | 0.074098 |
| CS3 | rs12339348 | 6233082 | 3.72E-30 | 0.073122 |
| CS3 | rs10815393 | 6240324 | 4.51E-30 | 0.070343 |

| | | | | |
|---|---|---|---|---|
| CS3 | rs7035413 | 6243119 | 6.89E-30 | 0.065363 |
| CS3 | rs7038893 | 6243392 | 9.09E-30 | 0.063107 |
| CS3 | rs17582919 | 6233376 | 7.05E-30 | 0.060414 |
| CS3 | rs17498196 | 6237547 | 9.47E-30 | 0.05379 |
| CS3 | rs10975504 | 6235009 | 9.17E-30 | 0.051897 |
| CS3 | rs72689561 | 6238750 | 1.54E-29 | 0.047636 |
| CS3 | rs78757963 | 6282511 | 2.44E-19 | 0.047079 |
| CS3 | rs10975507 | 6236977 | 1.65E-29 | 0.037797 |
| CS3 | rs76962799 | 6187132 | 1.40E-20 | 0.035635 |
| CS3 | rs7851246 | 6362365 | 7.51E-28 | 0.025082 |
| CS3 | rs7027505 | 6345740 | 4.08E-28 | 0.024066 |
| CS3 | rs16924428 | 6351111 | 6.52E-28 | 0.023837 |
| CS3 | rs10975547 | 6343945 | 4.60E-28 | 0.022612 |
| CS3 | rs59606381 | 6362124 | 1.38E-27 | 0.019674 |
| CS3 | rs7850988 | 6335760 | 6.74E-28 | 0.018757 |
| CS3 | rs72689565 | 6240953 | 2.84E-28 | 0.017263 |
| CS3 | rs10975558 | 6364449 | 2.35E-27 | 0.016619 |
| CS3 | rs16924356 | 6331610 | 7.02E-28 | 0.01507 |
| CS3 | rs10491836 | 6331421 | 7.55E-28 | 0.014958 |
| CS3 | rs2169287 | 6305904 | 2.05E-28 | 0.013794 |
| CS3 | rs10739094 | 6333685 | 1.60E-27 | 0.013582 |
| CS3 | rs72691711 | 6315489 | 2.72E-28 | 0.012367 |
| CS3 | rs112935616 | 6236830 | 5.44E-28 | 0.011728 |
| CS3 | rs7851000 | 6314487 | 2.86E-28 | 0.011704 |

**Supplementary table 3**. Comparison of *IL33* expression and protein levels between genotypes for SNPs rs10975479, rs1888909 and rs992969. Confidence interval (95% CI) values are shown for each group and genotype.

| **Dataset** | **SNP** | **95%CI** |
|---|---|---|
| Endobronchial brushing | rs1097547 | 0.11 |
| (mRNA) | rs1888909 | $7.2x10^{-3}$ |
| | rs992969 | 0.035 |
| Nasal epithelial cell brushings | rs10975479 | 0.023-0.380 |
| (mRNA) | rs1888909 | 0.144-0.355 |
| | rs992969 | 0.062-0.293 |
| Hutterites | rs10975479 | 0.87 |
| (protein) | rs1888909 | $2.6x10^{-3}$ |
| | rs992969 | $7.4x10^{-3}$ |
